# Identification and Characterization of Novel Outer Membrane Proteins of *Brachyspira pilosicoli*

**DOI:** 10.1101/2025.01.03.631275

**Authors:** Amisha Panda, Jahnvi Kapoor, B. Hareramadas, Ilmas Naqvi, Ravindresh Chhabra, Sanjiv Kumar, Anannya Bandyopadhyay

## Abstract

*Brachyspira pilosicoli* is a pathogenic, Gram-negative, spirochete bacterium that causes intestinal spirochetosis (IS) in birds, pigs, and humans and is distributed worldwide. This anaerobic intestinal bacterium colonizes the large intestine, potentially leading to colitis, diarrhoea, and decreased growth rate. Outer membrane proteins of Gram-negative bacteria play crucial roles in adhesion and host-pathogen interaction, helping the bacteria to evade the host immune system, and enhancing their virulence. However, *B. pilosicoli* outer membrane proteins are yet to be identified and characterized. Here, we report the computational discovery of 42 outer membrane β-barrel (OMBB) proteins in *B. pilosicoli* proteome predicted using a consensus-based computational framework. β-barrel architectures of the predicted proteins were confirmed by generating AlphaFold 3-based structural models. Structure-based functional annotation predicted putative functions for the identified OMBB proteins, including BamA homolog involved in folding and membrane insertion of OMPs, LptD homolog involved in transport of lipopolysaccharides into the OM, efflux pumps, transporters, enzymes, diffusion channels, and porins. Sequence variations across nine strains of *B. pilosicoli* were identified and mapped onto structural models, revealing that many of the variations were present on the surface exposed loop regions of the β-barrel structures. Our *in-silico* analysis has identified 42 OMBB proteins, including homologs of BamA, LptD, TolC, TonB-dependent receptors, CsgG. Seven of these were identified as hypothetical proteins. Computational characterization of the predicted OMBB proteins offer insights into their potential roles in physiology, virulence, and disease pathogenesis, highlighting their potential for applications in diagnostics, vaccine development, or therapeutic interventions.

## 1. Introduction

*Brachyspira pilosicoli,* previously known as *Serpulina pilosicoli*, is a zoonotic bacterium belonging to the Brachyspiraceae family within the order Spirochaetales of the phylum Spirochaetota.^1^ It is a Gram-negative, anaerobic, slow-growing, double-membraned, flagellated bacteria. *B. pilosicoli* causes intestinal spirochetosis (IS) in higher animals, including avian intestinal spirochetosis (AIS) in birds, porcine intestinal spirochetosis (PIS) in pigs and human intestinal spirochetosis (HIS) in humans. Spirochetal infections have been reported in the UK, continental Europe, Scandinavia, North America, Oceania, Iran, Malaysia, and South America.^2–7^ *B. pilosicoli* has a broad host range ^8,9^, including dogs, monkeys, water birds, game birds and humans.^8^ In IS, numerous brachyspiral cells penetrate the mucosal layer overlying enterocytes in the small intestine, attaching their one end to the luminal surface of the enterocytes, aided by surface lipoproteins. This attachment forms a distinctive layer resembling a ‘false brush border’.^8^ *B. pilosicoli* is the sole etiological agent of PIS, which is marked by diarrhoea and poor growth in pigs.^9,10^ AIS in chickens is associated with delayed onset of egg laying, production of wet and bloody faeces, reduced growth rate and diarrhoea.^8,11,3^ HIS is associated with various non-specific clinical symptoms, including abdominal pain, altered bowel patterns, chronic diarrhoea and rectal bleeding.^3,12–15^ Common risk factors associated with zoonotic transmission of *B. pilosicoli* to humans include exposure to faecal contaminated water, ^9^ ^16^ ^17,18^ rural or animal exposure, overcrowding, socioeconomic depression, travel to less developed countries, and being HIV positive or a homosexual male.^8^ AIS and PIS are under-reported diseases, bearing significant economic consequences for global food production. Although no comprehensive cost analysis for PIS exists, AIS alone costs the poultry industry approximately £18 million annually in the UK.^3^ Extrapolating from these figures, combined economic losses to both industries globally could reach approximately 1–2 billion USD annually.^19^ Antibiotic treatment is used for AIS, PIS, and HIS, but resistance has been reported.^18^ Antibiotics such as co-amoxicillin and metronidazole treat HIS, while pleuromutilins, macrolides and lincosamides are used for AIS and PIS.^18^ Although antibiotics are commonly used, no vaccines are currently available for preventing AIS or PIS, highlighting the urgent need for vaccine development.

*B. pilosicoli* strain 95/1000 has a single circular chromosome of approximately 2.59 Mb and lacks any extrachromosomal elements. The *B. pilosicoli* genome comprises 2,338 genes, with coding regions making up 85% of the total genome.^20^ Like other Gram-negative bacteria, *B. pilosicoli* consists of a central protoplasmic cylinder covered by a membrane sheath known as the outer membrane (OM).^21^ The exact composition of *B. pilosicoli* OM is not fully understood. Still, it is known to be extremely labile due to the high sterol content, which results in low resistance to osmotic stress and causes destabilisation when exposed to low ionic buffers.^22^ The *B. pilosicoli* outer envelope contains lipooligosaccharides (LOS) rather than lipopolysaccharides (LPS), which exhibits serological diversity across various strains.^23^ Bacteria with diderm envelopes possess a diverse family of outer membrane proteins (OMPs), characterized by β*-*barrel structures (OMBB) and lipooligosaccharides.^24,25^ β-barrels are protein structures made up of amphipathic anti-parallel β-strands that closes in on themselves, forming a cylindrical structure. β-barrels of OMPs are typically composed of an even number of β-strands, usually ranging from 8 to 36.^26^ β-strands are alternately connected on each side of the OM by long loops on the extracellular side and by shorter turns on the periplasmic side.^27^ OMBB proteins are involved in many functions, such as nutrient acquisition, membrane biogenesis, assembly of OMPs, adhesion, biofilm formation, efflux, proteolysis and even pilus formation.^28^ Thus, OMBB proteins represent a crucial area of research and a promising target for developing antibacterial therapies to combat pathogenic microbes. Notably, few OMPs of *B. pilosicoli* have been studied, BmpC (23 kDa lipoprotein)^22^, 45 kDa surface-exposed lipoprotein ^29^, and Bmp72.^30^ Christodoulides et al. employed an *in silico* reverse vaccinology approach to identify potential vaccine candidates from predicted OMBB proteins.^19^ Although few OMPs and lipoproteins of *B. pilosicoli* have been identified,^22,29–31^ identification and characterization of the complete OMPome are needed to define their potential role in disease pathogenesis, specifically to determine the extent of their involvement in critical processes such as attachment, virulence and generating host immune response.

The present study employed a comprehensive *in silico* approach to identify novel OMBB proteins in *B. pilosicoli.* A consensus of the output from OM localization prediction tools and β-barrel conformation prediction tools was considered for OMBB protein prediction. Through stringent screening criteria and manual curation, we selected 42 putative OMBB proteins. We generated deep-learning-based structural models of the proteins. Structural homologs were identified using the DALI server, unravelling the functional roles of the proteins. Amino acid sequence variations for the predicted proteins were obtained from nine strains of *B. pilosicoli* and were mapped on the structural models. Our study has identified 42 OMBB proteins of *B. pilosicoli*, computationally characterized their structure and function, and identified peptide regions crucial for bacterial pathogenesis.

## 2. Materials and Methods

### 2.1 Outer membrane protein prediction

Since reference genomes provide a streamlined, standardized, and taxonomically diverse representation of the RefSeq collection^32^, we selected the reference strain *B. pilosicoli* 95/1000, a porcine isolate, for our study. Using genome assembly ASM14372v1, the sequences of all proteins from *B. pilosicoli* 95/1000 genome were downloaded from NCBI.^32^

Peptide length, molecular weight, charge and isoelectric point (pI) for all the protein sequences were determined using the Pepstats tool from the EMBOSS package.^33^ Presence of signal peptide was determined using SignalP 5.0 and LipoP. SignalP was employed to predict signal peptide and its cleavage position in the protein sequences. SignalP server generates output for each protein sequence in the following formats: secretory signal peptide (Sec/SPI), lipoprotein signal peptide (Sec/SPII), Tat signal peptide (Tat/SPI), Tat lipoprotein signal peptide (Tat/SPII), Pilin signal peptide (Sec/SPIII) or the absence of any signal peptide (Other).^34^ LipoP server predicts lipoproteins of Gram-negative bacteria and discriminates between lipoprotein signal peptides, other signal peptides, and N-terminal transmembrane helices. The output is produced into four classes: secretory signal peptide (SpI), lipoprotein signal peptide (SpII), N-terminal transmembrane helix (TMH) and cytoplasmic protein (Cyt).^35^ The N-terminal TMH serves as an anchor, stabilizing the protein within the membrane. Hence, LipoP was used as a second tool to predict signal peptides. CELLO ^36^ and PSORTb ^37^ were utilized to predict the subcellular localization of proteins. Essential proteins from *B. pilosicoli* were predicted by searching for all proteins using BLASTP against the Database of Essential Genes (DEG v15.2).^38^ The DEG database is a repository of essential proteins from archaea, bacteria, and eukaryotes, and considers that proteins essential in one organism are likely to be essential in others. Specifically, proteins having an E-value of less than 1.0E-03 and a bit-score greater than 100 were considered.

The computational framework designed to select outer membrane proteins is detailed below and schematically represented in Figure 1. We employed a consensus-based computational approach to identify OMBB proteins where the output from four OMP prediction tools (one database-OMPdb ^39^, and three tools-MCMBB ^40^, TMBETADISC-RBF ^41^ and TMbed ^42^) was considered for OMP identification. Protein sequences were searched against the proteins in the OMPdb database to identify similar proteins (with E-value < 1.0E-03, bitscore > 100).

**Figure 1:**
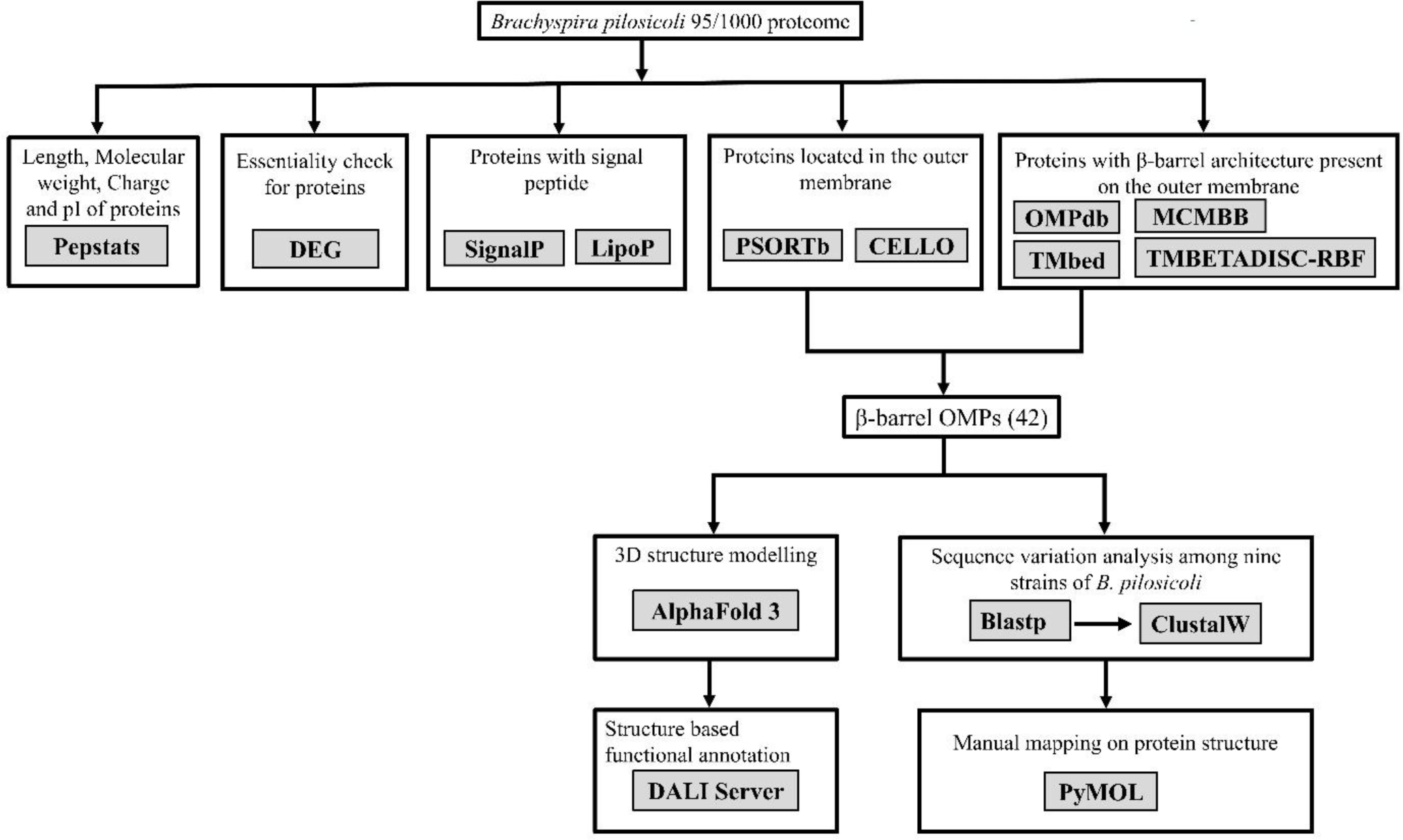
Computational framework for predicting OM-localized, β-barrel proteins from *Brachyspira pilosicoli* 95/1000. *B. pilosicoli* proteome was mined *in-silico* using various tools such as SignalP, LipoP, PSORTb, CELLO, OMPdb, MCMBB, Tmbed, TMBETADISC-RBF., Essentiality check for proteins was performed using DEG database. Size, charge, and isoelectric point of proteins were predicted using Pepstats. Structural models were generated using AlphaFold 3 server. These structural models were used as queries in the DALI server, and putative functions were annotated. Amino acid sequence variation analysis from nine strains of *B. pilosicoli* was performed using ClustalW and mapped onto structural models using the software PyMOL.

OMPdb is a database of integral β-barrel outer membrane proteins from Gram-negative bacteria. MCMBB discriminates β-barrel outer membrane proteins from globular proteins and α-helical membrane proteins. In MCMBB, a score greater than zero indicates that the protein is more likely to have a β-barrel conformation, whereas a score lower than zero shows that the protein is not a β-barrel. TMBETADISC-RBF server predicts OMPs based on radial basis function (RBF) network and position-specific scoring matrix (PSSM) profiles. TMbed, based on embeddings from protein language models (pLMs), predicts the propensity of each residue to form transmembrane helix (TMH), transmembrane β-strand (TMB), signal peptide, or other.

Using a consensus-based approach, outcomes obtained from the above OMP prediction tools were used to identify potential OMBB proteins. A final list of 42 OMBB proteins was compiled based on the number of tools predicting β-barrel architecture for each protein.

These proteins were categorized based on the number of tools providing positive results for a given protein sequence. This implied that higher confidence was assigned to those predicted as OMBB protein by more tools.

### 2.2 Structural modelling

Structural models of predicted OMBB proteins were generated using AlphaFold server, powered by AlphaFold 3.^43^ The modelling process incorporates physical and chemical constraints to accurately predict protein folding, resulting in atomic coordinates for each OMBB protein. Outputs of AlphaFold 3 include confidence metrices: pLDDT, PAE (predicted aligned error), and pTM and ipTM scores. The pTM (predicted template modelling) and ipTM (interface predicted template modelling) scores measure the accuracy of the entire structure.^44,45^ A pTM score above 0.5 and ipTM score above 0.8 indicate highly reliable predictions. The top ranked predictions based on pLDDT (predicted local distance difference test), were selected for figures and further analysis. The resulting atomic coordinate files were visualized using PyMOL.^46^ To validate the structures generated by AlphaFold 3, structural models were also generated using other modelling tools: ESMFold^47^, SWISS-MODEL^48^, RoseTTA^49^ and TrRosetta^50^.

### 2.3 Structure-based functional annotation using DALI server

Since many of the OM β-barrel proteins were unannotated hypothetical proteins, a structure-based approach was employed to unravel their functional roles. Atomic coordinates of the structural models were used as queries in the DALI server ^51^ with the full PDB search option, which ensures that the query will be compared against all protein structures available in the Protein Data Bank (PDB). The top-hit protein with the highest Z-score was selected for functional annotation. The Z-score is an optimized similarity score based on the sum of equivalent Cα-Cα distances between two proteins. A score above 20 confirms definite homology, 8–20 implies potential homology, while scores below 8 indicate insignificant similarity.^52^ Functions of the top-hit proteins were searched from the available literature to annotate functions to the predicted OMBB proteins.

### 2.4 Amino acid sequence variation among different strains of *B. pilosicoli*

Predicted OMBB proteins from the reference genome 95/1000 were searched for similar proteins (using BLASTP with E-value < 1.0E-03, bitscore > 100) in nine completed genomes of *B. pilosicoli* to analyze the amino acid sequence variations in the predicted β-barrel proteins (Table S2). Multiple Sequence Alignment (MSA) was done for orthologous sequences of each protein using ClustalW.^53^ Analysis of the MSA revealed amino acid variations among the orthologs. Mapping of amino acid sequence variations onto structural model was done using PyMOL.^46^

### 2.5 Structure alignment using US-Align server

Structural models were aligned using the US-align^54^ online web server to assess structural similarities and variations. Structural alignments were visualized, and figures were generated using PyMOL.

## 3. Results and Discussion

### 3.1 Prediction of OM β-barrel proteins using a consensus-based computational approach

A consensus-based computational framework was applied to the *B. pilosicoli* 95/1000 proteome, consisting of 2275 proteins, to identify OMBB proteins (Figure 1). Ten computational tools were used for predictions: Pepstats, DEG database, SignalP, LipoP, CELLO, PSORTb, OMPdb, MCMBB, TMBETADISC-RBF, and TMbed (Table S1).

Prediction outputs from all the tools were combined for each protein. Tools that specifically predict OMBB proteins (OMPdb, MCMBB, TMBETADISC-RBF, and TMbed) were utilized for OMBB protein prediction (Table S1). Through stringent screening criteria and manual curation, 42 OMBB proteins were selected and subsequently classified into two groups: Group A, with 13 proteins predicted as OMBB by all four tools, and Group B, with 29 proteins predicted as OMBB by any three tools (Table 1).

**Table 1:**
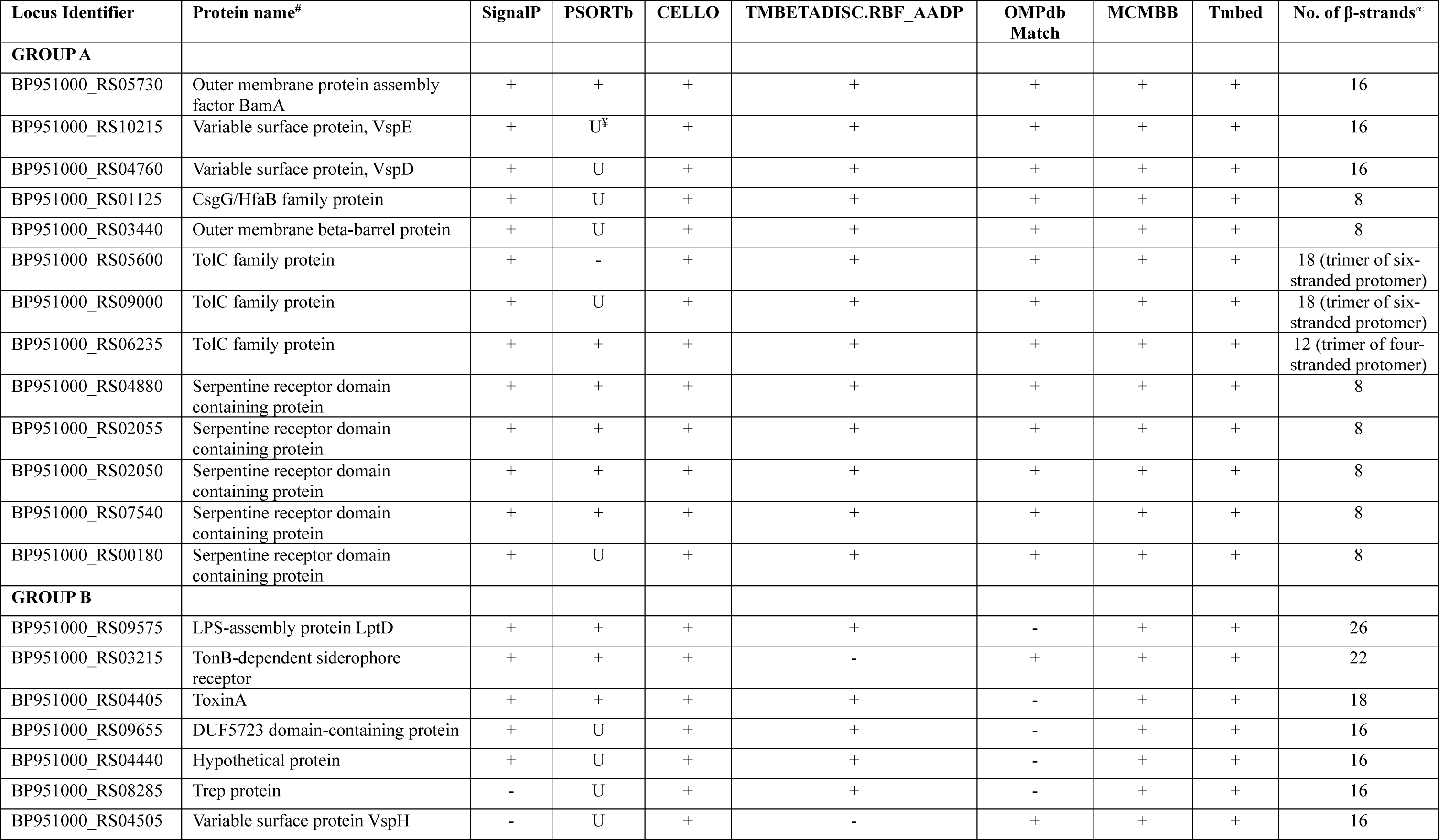
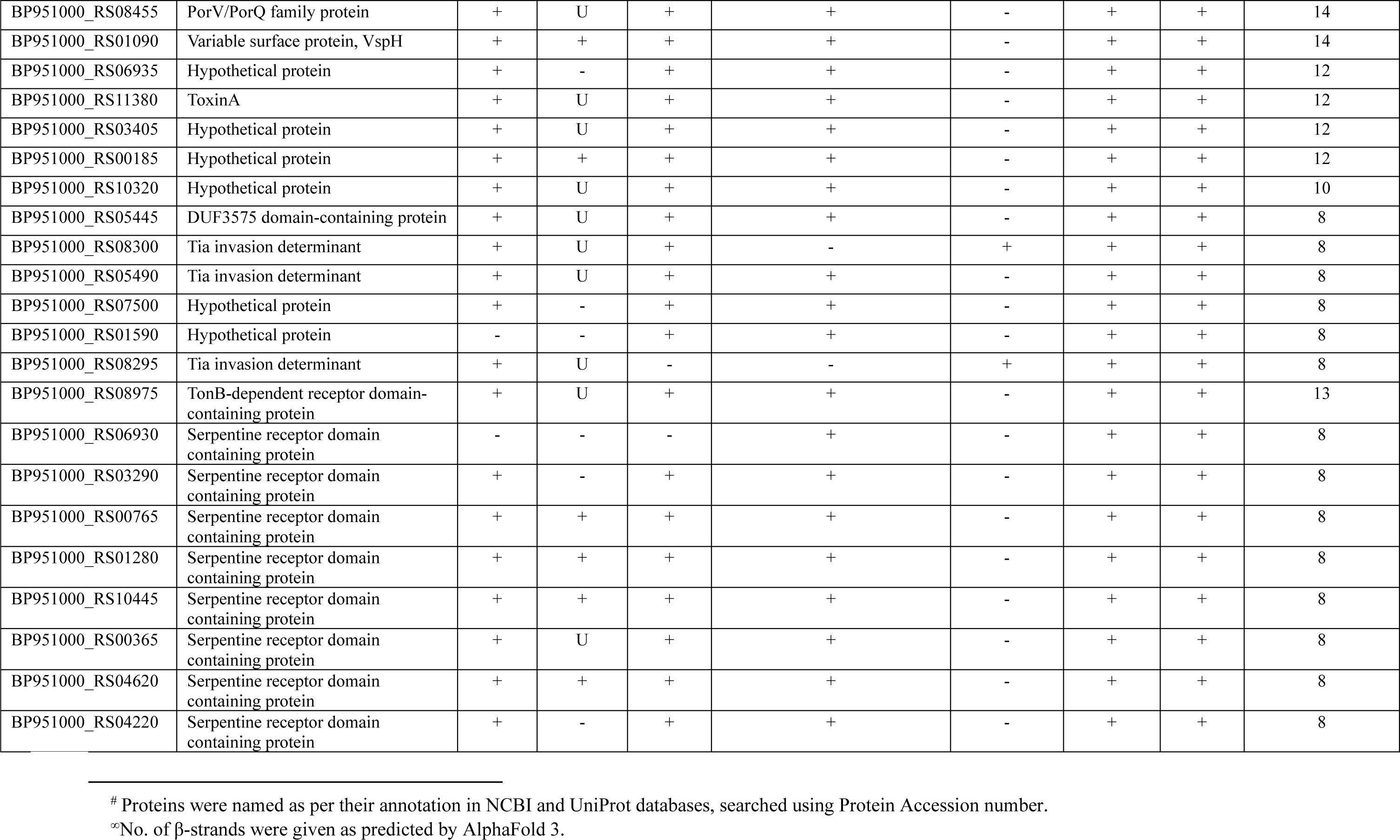
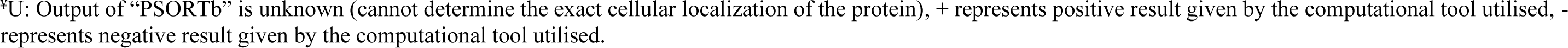
Predicted outer membrane β-barrel proteins from *B. pilosicoli* 95/1000.

To gain structural insights into the predicted proteins, we searched for their structures in the Protein Data Bank (PDB) but found no available structures. Using AlphaFold 3^43^, structural models of the predicted proteins were generated, revealing typical features of OM proteins, such as β-barrel architectures with central pores, periplasmic loops, and surface-exposed loops. Since most proteins were unannotated, a structure-based approach using the DALI protein structure comparison server was employed to annotate their putative functions. Top hits with the highest Z-score were considered for functional annotation. This analysis revealed structural homologs of well-known proteins, including BamA, LptD, NspA, OmpF, OMPLA, BtuB, providing valuable insights into their possible roles (Table 2). OM proteins are positioned on the bacterial surface, serving as the primary interface between host and pathogen. Since OMPs are subjected to strong selection pressures to adapt to the host environment, monitoring sequence variation in these proteins is crucial for understanding the evolutionary force acting on the pathogen (Table 3).^55^ Towards this, we analyzed the amino acid sequences of the predicted OMBB proteins for residues exhibiting sequence variation across nine *B. pilosicoli* strains (Table S2). Mapping the variations onto the structural models revealed that many variations were located on extracellular loops, which tend to interact with the host environment (Table 3).

**Table 2:**
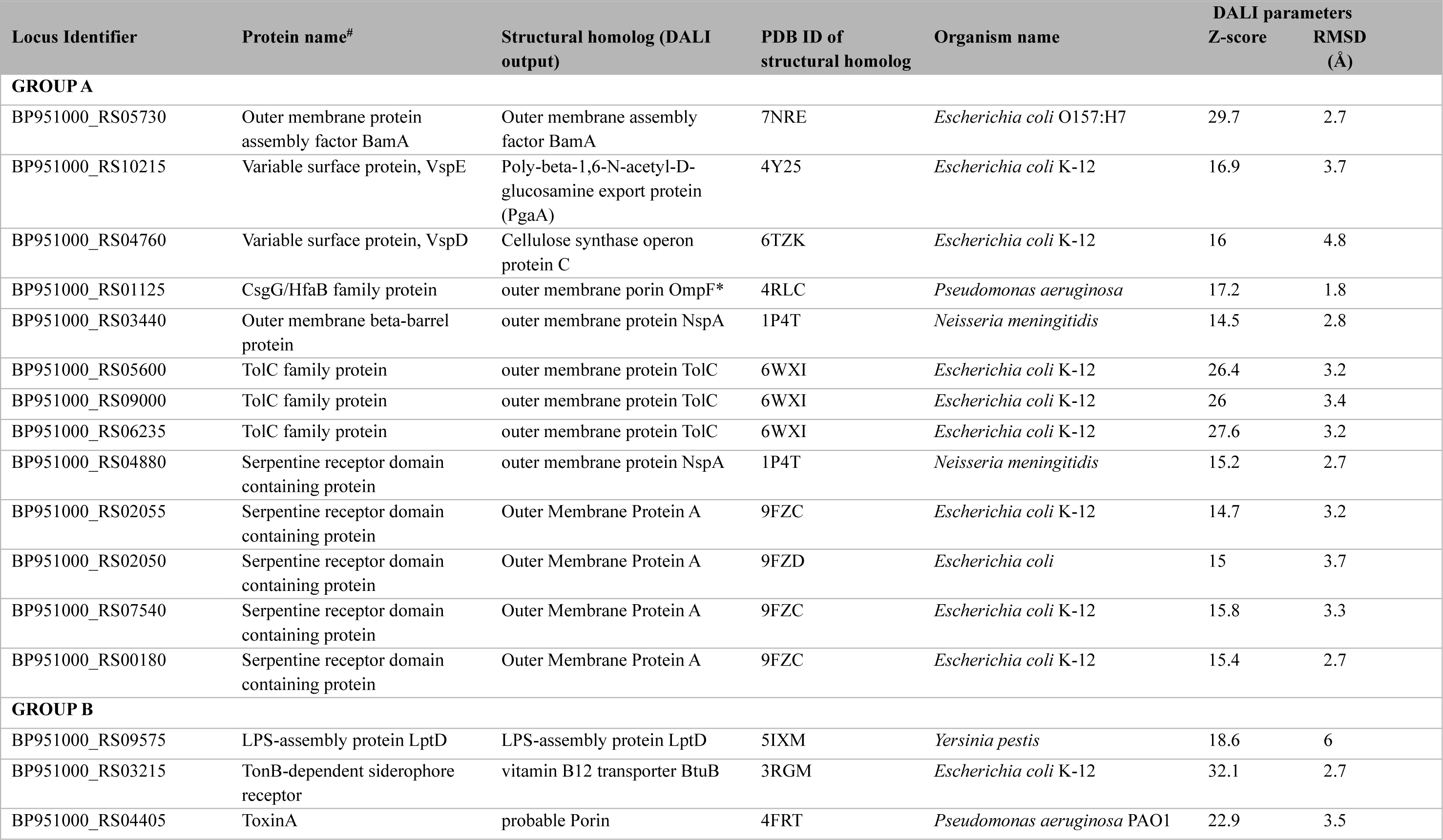
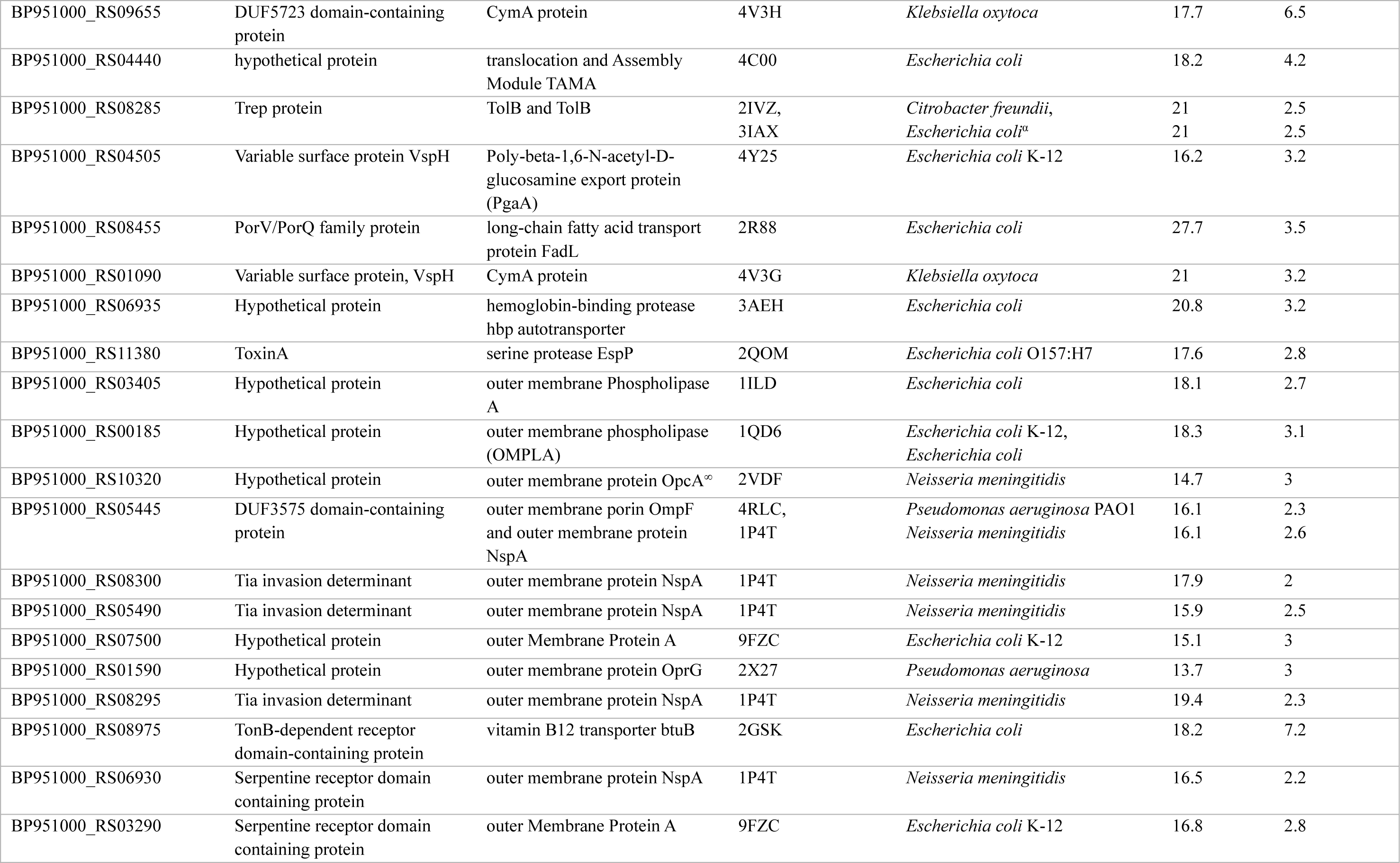
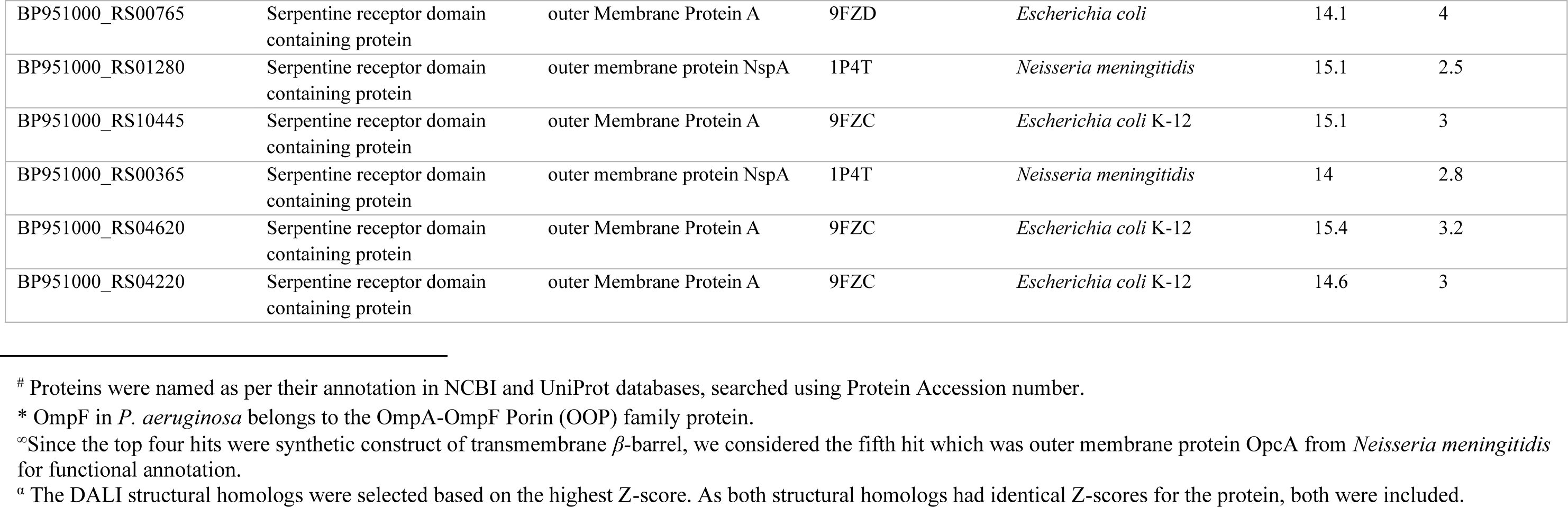
Closest PDB structures matched through DALI Server based on the highest Z-score for the predicted β-barrel OMPs.

**Table 3:** Sequence variations among nine strains: Total variations and variations in the extracellular loops of predicted OM β-barrel proteins.

| Locus Identifier | Protein name<sup>#</sup> | Total variations | Variations present on predicted ECL region |
| --- | --- | --- | --- |
| **GROUP A** |  |  |  |
| BP951000_RS05730 | Outer membrane protein assembly factor BamA | D60, A184, V465, A467, F512 | None |
| BP951000_RS10215<sup>n</sup> | Variable surface protein, VspE | 208 variations | Not determined<sup>u</sup> |
| BP951000_RS04760<sup>n</sup> | Variable surface protein, VspD | 260 variations | Not determined<sup>u</sup> |
| BP951000_RS01125 | CsgG/HfaB family protein | S63, D79, T190, I210, L380 | None |
| BP951000_RS03440 | Outer membrane beta-barrel protein | F24, V47, V64, N110, D169, A197 | V47 |
| BP951000_RS05600 | TolC family protein | T246, N499 | None |
| BP951000_RS09000 | TolC family protein | S90, S131 | S90 |
| BP951000_RS06235 | TolC family protein | K2, N3, F5, V6, F7, I8, I10, L12, S16, S25, N33, I42, E43, L93, S105, E136, I137, T210 | L93 |
| BP951000_RS04880 | Serpentine receptor domain containing protein | N69 | None |
| BP951000_RS02055 | Serpentine receptor domain containing protein | H101, N163, M235 | N163, M235 |
| BP951000_RS02050 | Serpentine receptor domain containing protein | A27, L28, T108, A228, I247 | T108, A228 |
| BP951000_RS07540 | Serpentine receptor domain containing protein | M1, K2, K3, I4, I5, L6 | None |
| BP951000_RS00180 | Serpentine receptor domain containing protein | I64, M115 | None |
| **GROUP B** |  |  |  |
| BP951000_RS09575 | LPS-assembly protein LptD | N14, G137, I257, I382, E454, D600, G944 | D600 |
| BP951000_RS03215<sup>n</sup> | TonB-dependent siderophore receptor | 59 variations | Not determined<sup>u</sup> |
| BP951000_RS04405 | ToxinA | M1, H2, R3, I4, I6, L8, T9, M18, V19, T24, N32, S34, N41, F84, K90, N98, I101, S102, N104, S175, Q183, I264, T303 | N41, F84, K90, N98, S175, Q183 |
| BP951000_RS09655<sup>n</sup> | DUF5723 domain-containing protein | 315 variations | Not determined<sup>u</sup> |
| BP951000_RS04440 | hypothetical protein | K104, K113, S117, Y124, I132, T134, N151, G153, L243, V252, L254, S308, N321 | K104, L243 |
| BP951000_RS08285<sup>n</sup> | Trep protein | 43 variations | Not determined<sup>u</sup> |
| BP951000_RS04505<sup>n</sup> | Variable surface protein VspH | 247 variations | Not determined<sup>u</sup> |
| BP951000_RS08455 | PorV/PorQ family protein | L12, S20, N22, A117, R187, S253 | N22, A117, |
| BP951000_RS01090 | Variable surface protein, VspH | E258 | E258 |
| BP951000_RS06935 | Hypothetical protein | S9, I10, V13, R298 | None |
| BP951000_RS11380 | ToxinA | M126, M154, I278 | M126 |
| BP951000_RS03405 | Hypothetical protein | M1, R2, L3, K4, F5, F6, F7, L8, I9, F10, L11, F12, L13, S14, L15, S16, L17, Y18, | None |
|  |  | T19, Q20, D21, N22, E23, A24 |  |
| BP951000_RS00185<sup>η</sup> | Hypothetical protein | 58 variations | Not determined<sup>μ</sup> |
| BP951000_RS10320 | Hypothetical protein | L18, D48, E55, F251, G285, E400, Y401, G402, I403, F404, T405, K406, Q407, L408, A409, I410, S411, F412, I413, P414, I415, N416, I417, R418, F419 | None |
| BP951000_RS05445 | DUF3575 domain-containing protein | K2, I7, A79, N87, H89, K158 | N87, H89 |
| BP951000_RS08300 | Tia invasion determinant | L143, N156, S200 | None |
| BP951000_RS05490<sup>η</sup> | Tia invasion determinant | 61 variations | Not determined<sup>μ</sup> |
| BP951000_RS07500<sup>η</sup> | Hypothetical protein | 183 variations | Not determined<sup>μ</sup> |
| BP951000_RS01590 | Hypothetical protein | V205, I215, V221 | None |
| BP951000_RS08295 | Tia invasion determinant | N34, I49, V123, S141, I144, V167 | N34 |
| BP951000_RS08975 | TonB-dependent receptor domain-containing protein | D32, T371 | D32 |
| BP951000_RS06930<sup>η</sup> | Serpentine receptor domain containing protein | 135 variations | Not determined<sup>μ</sup> |
| BP951000_RS03290 | Serpentine receptor domain containing protein | None | None |
| BP951000_RS00765<sup>η</sup> | Serpentine receptor domain containing protein | 70 variations | Not determined<sup>μ</sup> |
| BP951000_RS01280 | Serpentine receptor domain containing protein | V32, A83, V124, E210, T237 | E210 |
| BP951000_RS10445 | Serpentine receptor domain containing protein | G228 | None |
| BP951000_RS00365 | Serpentine receptor domain containing protein | V72, Q77, I84, D156, D159, V168, N177, A200, T216, I222, Y226 | D156, D159, A200 |
| BP951000_RS04620 | Serpentine receptor domain containing protein | K2, E95, A140, V194 | None |
| BP951000_RS04220<sup>η</sup> | Serpentine receptor domain containing protein | 218 variations | Not determined<sup>μ</sup> |

### 3.2 Identification of OM Proteins Group A

Group A comprised of 13 proteins, consisting of three proteins with 16 stranded β-barrel domain (BP951000_RS05730, BP951000_RS10215, BP951000_RS04760); seven proteins with eight-stranded β-barrel domain (BP951000_RS02055, BP951000_RS02055, BP951000_RS07540, BP951000_RS01125, BP951000_RS00180, BP951000_RS03440, BP951000_RS04880), and three TolC family proteins (BP951000_RS05600, BP951000_RS09000, BP951000_RS06235) (Table 1). Out of the seven eight-stranded β-barrel proteins, five (BP951000_RS02055, BP951000_RS02055, BP951000_RS07540, BP951000_RS00180 and BP951000_RS04880) are annotated as serpentine receptor domain containing proteins.

#### 3.2.1 BP951000_RS05730, BamA

BP951000_RS05730 is annotated as BamA in *B. pilosicoli* strain 95/1000. Bacterial BamA, along with BamBCDE, forms *<u>β</u>*-barrel <u>a</u>ssembly <u>m</u>achine (BAM) complex which is involved in assembly and insertion of β-barrel proteins into the OM.^56^ BP951000_RS05730 has been identified as an essential protein in DEG database. BP951000_RS05730 structural model, generated using AlphaFold 3, showed a characteristic BamA bipartite structure consisting of a periplasmic N-terminal region and a C-terminal β-barrel domain (Figure 2A). The N-terminal segment contains five <u>po</u>lypeptide <u>tr</u>ansport-<u>a</u>ssociated (POTRA) domains (P1–P5) comprising characteristic β1-α1-α2-β2-β3 motif. In other well-characterized BamA proteins, these domains form a scaffold for binding of BamBCDE proteins and facilitate the folding of OMPs.^57^ *B. pilosicoli* BamA consists of 16 antiparallel β-strands, with a characteristic lateral gate between strands 1 and 16. Structural-homology search using DALI server revealed the closest match (Z-score = 29.7 and RMSD = 2.7 Å) with BamA of *E. coli* O157:H7 (PDB ID: 7NRE) (Table 2). This result validated the functional annotation of BamA in *B. pilosicoli*.

**Figure 2:**
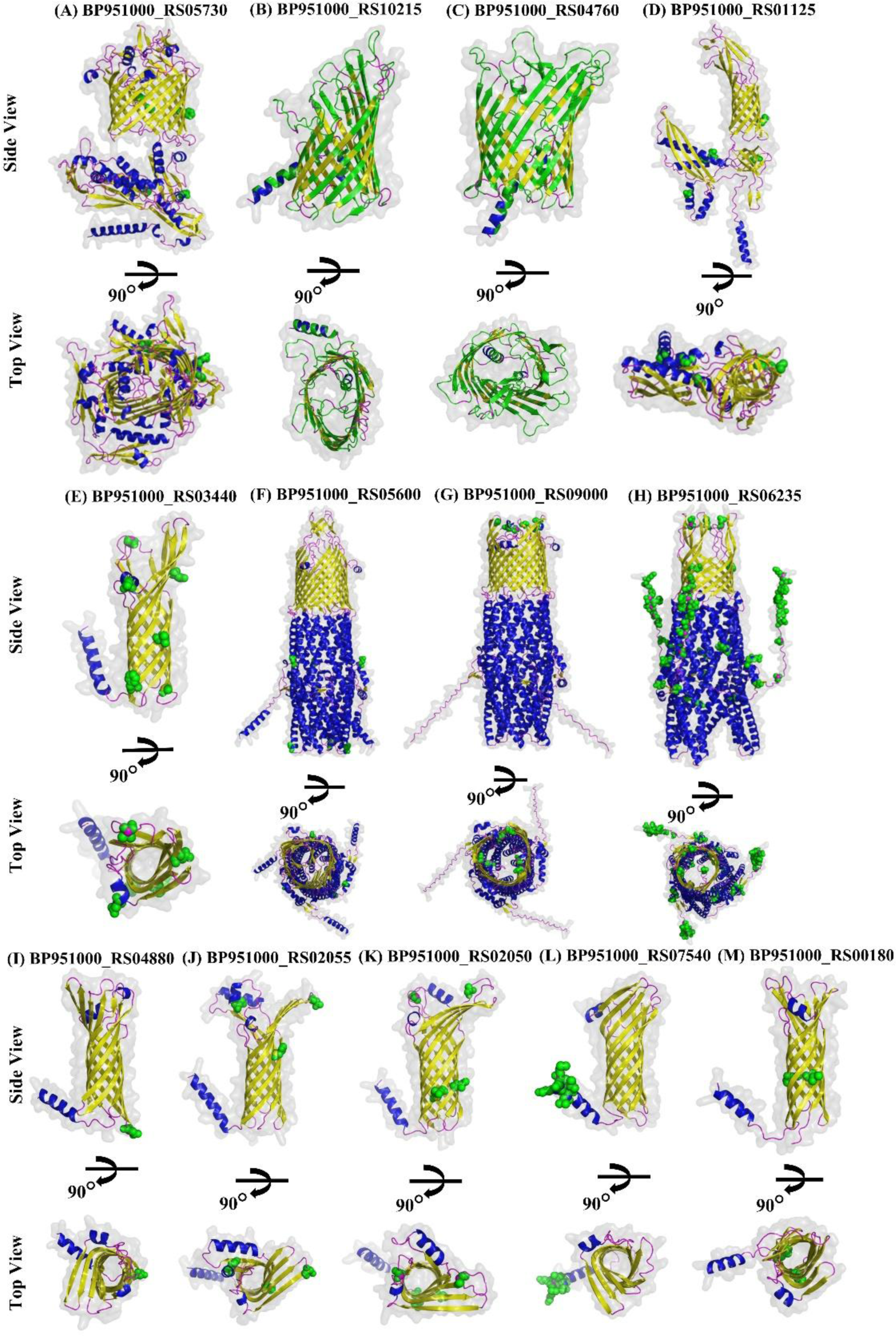
Structural models of β-barrel outer membrane proteins in Group A. Group A consists of 13 proteins. (A), (B) and (C) comprise 16 β*-*strands. (F), (G) and (H) depict predicted trimeric structures of TolC family proteins. The remaining proteins have eight-stranded β-barrel structures. β-strands, α-helices and loops are shown in yellow, blue, and magenta colors, respectively. Green spheres indicate amino acid variations across nine strains of *B. pilosicoli*. Proteins exhibiting more than 40 variations are represented using ribbon representation in green colour.

Sequence comparison of BP951000_RS05730 across nine strains of *B. pilosicoli* revealed five variations (D60, A184, V465, A467 and F512) (Tables-3, S3). When mapped on the structural model, V465, A467, and F512 were present in the β-barrel transmembrane (TM) domain, while D60, V184 were located in the periplasmic region of the protein (Tables-3, S3).

#### 3.2.2 BP951000_RS10215

BP951000_RS10215 is annotated as hypothetical protein in NCBI. However, it is annotated as variable surface protein, VspE in the UniProt database. Variable surface proteins (Vsps) are OMPs identified so far in *Brachyspira hyodysenteriae* and *Mycoplasma bovis,* and are used by these pathogenic bacteria to adapt to host conditions and enhance colonization.^58,59^ These proteins can show reversible on/off switching of their expression or undergo antigenic changes by expressing alternative protein phenotypes.^58^ They can function as mediators for bacterial attachment to host cells.^60^ Vsp-like proteins have been identified in *B. hyodysenteriae*, *B. pilosicoli* and *Mycoplasma*,^29,58,61^ however homologs in other bacteria remain undiscovered, highlighting their unique role in these pathogens.

*B. pilosicoli* VspE, BP951000_RS10215 has a secretory signal peptide with a cleavage site between residues 21 and 22. The structural model generated by AlphaFold 3 revealed a β-barrel architecture consisting of 16 β-strands, with the ninth and tenth strands longer than the others, giving an elliptical shape to the extracellular surface of the barrel (Figure 2B). BP951000_RS10215 showed the best structural alignment with the C-terminal β-barrel domain of Poly-beta-1,6-N-acetyl-D-glucosamine export protein (PgaA) of *E. coli* K-12 (PDB ID: 4Y25) (Table 2), with an RMSD value of 3.7 Å and a Z-score of 16.9. *E. coli* PgaA contains a 16-stranded β-barrel domain at the C-terminal and eight periplasmic tetratricopeptide repeats (TPRs) at the N terminal. In contrast, BP951000_RS10215 lacks periplasmic TPR domains. PgaA facilitates the translocation of PNAG (Poly-beta-1,6-N-acetyl-D-glucosamine) polymer from the periplasm to the cell surface, a key step in biofilm formation.^62–64^ The structural similarity of β-barrels between *E. coli* PgaA and BP951000_RS10215 implies a possible role of the Brachyspiral protein in translocation. Given that it shows high sequence and structural homology with *B. hyodysenteriae,* there is a high probability that BP951000_RS10215 plays a role in adherence and host colonization.^58,59,60^ Sequence variation of BP951000_RS10215 across nine strains of *B. pilosicoli* revealed 208 variations and several deletions (Table S4). These variations were spread throughout the protein (Figure 2B). This high degree of variability suggests the ability of a bacteria to adapt rapidly to changing environments or host immune responses.^58^

#### 3.2.3 BP951000_RS04760

BP951000_RS04760 is annotated as variable surface protein, VspD in UniProt database. However, it is annotated as variable surface family protein in NCBI. As discussed in section 3.2.2, Vsp proteins are involved in bacterial attachment to host cells.^60^ *B. hyodysenteriae* VspD is a virulence factor and a potential vaccine development target.^65^ BP951000_RS04760 carries a secretory signal peptide with a cleavage position between residues 21 and 22. The structural model revealed a 16-stranded β-barrel architecture, with varying strand sizes creating an elliptical barrel surface on the extracellular side (Figure 2C). The protein showed the closest structural match (Z-score = 16 and RMSD = 4.8 Å) with β-barrel domain of cellulose synthase operon protein C (porin BcsC) (PDB ID: 6TZK) of *E. coli* K-12 (Table 2). BcsC is a 16-stranded β-barrel protein with a periplasmic domain that consists of 19 TPRs (Tetratricopeptide repeats), facilitating the secretion of phosphoethanolamine-cellulose (pEtN-cellulose) across the outer membrane.^66–70^ In contrast, BP951000_RS04760 lacks TPRs. The structural homology between β-barrels of *E. coli* BcsC and BP951000_RS04760 implies the possible role of BP951000_RS04760 in translocation, including bacterial adhesion and virulence similar to VspD of *B. hyodysenteriae*. Among nine strains of *B. pilosicoli*, BP951000_RS04760 revealed variations at 260 positions and deletions at multiple positions, indicating its high variability (Table S4). These variations were spread throughout the protein (Figure 2C).

#### 3.2.4 BP951000_RS01125

BP951000_RS01125 is annotated as Curli production assembly transport component, CsgG in *B. pilosicoli*. The CsgG curli production assembly transport component OMP is essential for the secretion of curli, functional amyloid fibers that form the main protein component of biofilm extracellular matrices in Bacteroidetes and Proteobacteria, playing key roles in pathogenesis.^71^ Curli fimbriae are involved in the initial colonization of the host, as well as in bacterial persistence and invasion.^72–74^ SignalP predicted a lipoprotein signal peptide with a cleavage position between residues 14 and 15. LipoP predicted it as a cytoplasmic protein.

Structural model of the protein showed a β-barrel architecture comprising eight β-strands extending into the periplasm via periplasmic domain (Figure 2D). DALI server results showed that BP951000_RS01125 exhibited best match with OmpF (PDB ID: 4RLC) of *Pseudomonas aeruginosa* (Table 2), with a Z-score of 17.2 and an RMSD value of 1.8 Å. It has been suggested that OmpF, eight-stranded β-barrel protein, is involved in biofilm formation, production of outer membrane vesicles, adhesion, and host immune system modulation.^75–81^ It can be inferred that BP951000_RS01125 might serve similar functions to OmpF of *P. aeruginosa*. Among the nine strains of *B. pilosicoli*, BP951000_RS01125 showed variations at five positions (S63, D79, T190, I210, L380) (Tables-3, S3). When mapped onto the structural model, one variation (L380) was found to be present in the β-barrel domain while rest of the variations were found to be present in the periplasmic region of the protein (Table S3).

#### 3.2.5 BP951000_RS03440

BP951000_RS03440, annotated as hypothetical protein in UniProt database, has a secretory signal peptide with a cleavage position between residues 21 and 22. However, it is annotated as Outer membrane β-barrel protein in NCBI. Structural model generated by AlphaFold 3 revealed a β-barrel architecture comprising eight β-strands, four extracellular loops, and three periplasmic loops (Figure 2E). BP951000_RS03440 showed best structural alignment with NspA (PDB ID: 1P4T) of *Neisseria meningitidis* with the highest Z-score of 14.5 (RMSD = 2.8 Å) (Table 2). NspA is an eight-stranded β-barrel protein, playing an important role in attachment of the bacteria to the host immune system and promotes their colonization. It has been predicted as a potential vaccine candidate. ^82–85^ It is plausible to speculate that BP951000_RS03440 might be involved in attachment of bacteria to the host immune system. Multiple Sequence Alignment of BP951000_RS03440 homologs across nine strains of *B. pilosicoli* showed six variations (F24, V47, V64, N110, D169, A197) (Tables-3, S3). When mapped onto the structural model, one variation (V47) was found to be present in the ECL region. Four variations (F24, V64, N110, A197) were found to be present in the TM region and one was located in the extracellular region of the protein (Table S3).

#### 3.2.6 BP951000_RS05600

BP951000_RS05600 has been annotated as putative outer membrane component of multidrug efflux system in UniProt database. However, it has been annotated as TolC family protein in NCBI. Both SignalP and LipoP identified a secretory signal peptide in this protein. Structural model generated by AlphaFold 3 was a trimer forming 18-stranded β-barrel where each subunit (protomer) comprised of six β-strands forming approximately one-third of the barrel structure (Figure 2F). Since no 18-stranded β-barrel TolC structures were reported in the literature, we validated the structural model generated by AlphaFold 3 using other modelling tools, including TrRosetta, RoseTTAFold, ESMFold, and SWISS-MODEL. The structural models showed high similarity and aligned well with the AlphaFold 3 prediction. Structural alignment of the five monomeric BP951000_RS05600 models constructed using five different tools yielded an RMSD of 3.96 Å, confirming model accuracy (Figure S1). Like the canonical TolC, the TM barrel extends into the periplasm as an α-helical structure, forming a periplasmic tunnel connected to the OM (Figure 2F)..^86^ The periplasmic entry to the tunnel is blocked, likely to prevent leakage through the OM, as the β-barrel domain remains constantly open.^87^ BP951000_RS05600 showed the best structural alignment with TolC (PDB ID: 6WXI) of *E. coli* K-12 (RMSD = 3.2 Å and Z-score = 26.4) (Table 2). TolC is involved in secretion of hemolysin ^88,89^, colicin import ^90,91^, efflux of antibiotics ^92^ and act as cell surface receptor for bacteriophage.^93^ Therefore, it can be speculated that BP951000_RS05600 might have functions similar to TolC of *E. coli* K-12. Sequence comparison across nine strains of *B. pilosicoli* showed two variations (T246, N499) (Tables-3, S3). Mapping the variations onto the structural model revealed that all the variations were present in the periplasmic α-helical regions of the protein (Table S3).

#### 3.2.7 BP951000_RS09000

BP951000_RS09000 has been annotated as an outer membrane efflux protein in UniProt database. However, it has been annotated as TolC family protein in NCBI. It carries secretory signal peptide. Structural model revealed a trimeric structure consisting an 18-stranded β-barrel where each subunit (protomer) comprised of six β-strands forming approximately one-third of the barrel structure (Figure 2G). Similar to BP951000_RS05600 (section 3.2.6), we validated the structural model generated by AlphaFold 3 using other modelling tools. Further, we structurally aligned the monomeric structures of BP951000_RS09000 generated by all five tools, resulting in an RMSD value of 3.85 Å (Figure S2). This suggested that the models were very similar and consistent with AlphaFold 3 prediction. Like canonical TolC, the barrel extends into the periplasm as an α-helical structure, creating a periplasmic tunnel within its lumen that connects to the OM (Figure 2G).^86^ The periplasmic entry to the tunnel is blocked, likely preventing leakage through the OM, as the β-barrel domain remains continuously open.^87^ BP951000_RS09000 exhibited structural match with TolC (PDB ID: 6WXI) of *E. coli* K-12 (Table 2), corresponding to a Z-score of 26 (RMSD = 3.4 Å). As described previously in section 3.2.6, BP951000_RS09000, structural homolog of TolC of *E. coli* K-12, might be involved in secretion of hemolysin ^88,89^, colicin import ^90,91^, efflux of antibiotics ^92^ and act as cell surface receptor for bacteriophage.^93^ Amino acid sequence variation across nine strains of *B. pilosicoli* showed two variations (S90, S131) (Tables-3, S3). When mapped onto the structural model, S90 was present in the ECL region (Figure 2G) while S131 was present in the TM β-barrel domain of the protein (Table S3).

#### 3.2.8 BP951000_RS06235

BP951000_RS06235 is annotated as a TolC family protein in the UniProt database. AlphaFold 3-generated structural model was a trimer forming 12-stranded β-barrel where each subunit comprised of four β-strands forming approximately one-third of the barrel structure (Figure 2H). As described in section 3.2.6, we generated structural models using other modelling tools (ESMFold, SWISS-MODEL, RoseTTA and TrRosetta) and performed a structural alignment of the monomeric structure of BP951000_RS06235 generated by all five tools, which resulted in an RMSD value of 3.26 Å (Figure S3). The barrel extends into the periplasm as an α-helical structure, forming a periplasmic tunnel within its lumen that links to the OM (Figure 2H).^86^ The periplasmic entry to the tunnel is blocked, likely to prevent leakage through the OM, as the β-barrel domain remains constantly open.^87^ BP951000_RS06235 showed the best structural match with TolC (PDB ID: 6WXI) of *E. coli* K-12 (Table 2) with a Z-score of 27.6 and an RMSD value of 3.2 Å. Considering *E. coli* K-12 TolC’s role (discussed in section 3.2.6), BP951000_RS06235 may serve a similar function. 18 amino acid residues were found to be varying across the nine strains of *B. pilosicoli*: K2, N3, F5, V6, F7, I8, I10, L12, S16, S25, N33, I42, E43, L93, S105, E136, I137, T210 (Tables-3, S3). When mapped onto the structural model, L93 in the ECL region (Figure 2H), S105 in the β-barrel region and rest of the variations were present in the periplasmic region of the protein (Table S3).

#### 3.2.9 Eight-stranded β-barrel proteins annotated as Serpentine receptor domain containing proteins

A total of thirteen proteins, annotated in the UniProt database as serpentine receptor domain containing proteins, were predicted as outer membrane β-barrel proteins in our study. Five of these proteins-BP951000_RS02055, BP951000_RS02050, BP951000_RS07540, BP951000_RS00180 and BP951000_RS04880, were categorized under Group A (predicted as OMBB proteins by all tools). These proteins are annotated as hypothetical in NCBI, except BP951000_RS02055, identified as an outer membrane β-barrel protein. Serpentine receptors (SRs), also known as G-Protein Coupled Receptors (GPCRs), are membrane-bound, heptahelical proteins found exclusively in eukaryotes, crucial for mediating cell-environment communication.^94,95^ The *B. pilosicoli* proteins with this annotation were predicted to contain a secretory signal peptide. Structural models from AlphaFold 3 revealed β-barrel conformations with eight β-strands, four extracellular loops, and three periplasmic loops (Figure 2I–M). These structures were validated by additional tools (ESMFold, SWISS-MODEL, RoseTTAFold and TrRosetta). Since GPCRs are absent in prokaryotes and structural models generated using five different tools predicted TM β-barrels rather than α-helices, we conclude that these proteins are misannotated as serpentine receptor domain-containing proteins in the UniProt database. BP951000_RS02055, BP951000_RS07540 and BP951000_RS00180 showed their best structural alignment with the N-terminal β-barrel domain of *E. coli* K-12 OmpA (PDB ID: 9FZC) (Table 2). BP951000_RS02050 exhibited the best structural match with *E. coli* OmpA β-barrel domain (PDB ID: 9FZD) (Table 2). Unlike *E. coli* OmpA, which consists of an N-terminal β-barrel domain and a periplasmic C-terminal domain, BP951000_RS02055, BP951000_RS02050, BP951000_RS07540, and BP951000_RS00180 lack the periplasmic domain, highlighting a significant structural difference. *E. coli* OmpA functions as a receptor for bacteriophages, facilitates colicin vesicles action, mediates F-dependent conjugation, maintains membrane integrity, and serves as a diffusion channel for small solutes.^96–103^ It also contributes to virulence and pathogenicity of *E. coli*, making it a key target in the immune response.^104^ Due to high structural homology of BP951000_RS02055, BP951000_RS02050, BP951000_RS07540 and BP951000_RS00180 with *E. coli* OmpA, they are likely to be involved in similar roles. BP951000_RS04880 showed the best structural alignment with NspA (PDB ID: 1P4T) of *N. meningitidis* (Z-score= 15.2, RMSD=2.7 Å) (Table 2). NspA is an eight-stranded β-barrel protein that plays an important role in the attachment of the bacteria to the host immune system and promotes their colonization. It has been predicted as a potential vaccine candidate.^82–85^ This suggests that BP951000_RS04880, based on its high structural homology with NspA, might have a role in adhesion and may be explored experimentally as a potential candidate for vaccine development. Amino acid sequence variation analysis for all five proteins across nine strains of *B. pilosicoli* revealed several variations (Tables 3 and S3). BP951000_RS02055 and BP951000_RS02050 revealed variations in the ECL regions (Figure 2J, 2K). Variations in the rest of the proteins were present in the TM and/or periplasmic regions (Figure 2I, 2L, 2M; Tables 3 and S3).

##### Group B

Group B comprised of 29 proteins: 26-stranded β-barrel protein (BP951000_RS09575); 22-stranded β-barrel protein (BP951000_RS03215); 18-stranded β-barrel protein (BP951000_RS04405); 16-stranded β-barrel proteins (BP951000_RS09655, BP951000_RS04440, BP951000_RS08285, BP951000_RS04505); 14-stranded β-barrel proteins (BP951000_RS08455, BP951000_RS01090); 13-stranded β-barrel protein(BP951000_RS08975); 12-stranded β-barrel proteins (BP951000_RS06935, BP951000_RS11380, BP951000_RS03405, BP951000_RS00185); 10-stranded β-barrel protein (BP951000_RS10320); 8-stranded β-barrel proteins (BP951000_RS05445, BP951000_RS08300, BP951000_RS05490, BP951000_RS07500, BP951000_RS01590, BP951000_RS08295); 8-stranded β-barrel Serpentine receptor domain containing proteins (BP951000_RS06930, BP951000_RS03290, BP951000_RS00765, BP951000_RS01280, BP951000_RS10445, BP951000_RS00365, BP951000_RS04620, BP951000_RS04220) (Table 1).

#### 3.2.10 BP951000_RS09575 (LptD)

BP951000_RS09575 has been annotated as LptD in *B.pilosicoli*.^20^ LptD is a part of the lipopolysaccharide transport (LPT) system, which transports lipopolysaccharides (LPS) from the inner leaflet of OM to the cell surface in Gram-negative bacteria.^105^ BP951000_RS09575 has a secretory signal peptide with a cleavage position between residues 19 and 20. It has been identified as an essential protein in DEG database. The structural model of BP951000_RS09575 revealed a large β-barrel architecture consisting of an antiparallel arrangement of 26 β-strands, forming a cylindrical structure that spans the OM (Figure 3A). It has a lateral opening between strands 1 and 26. Like other LptD orthologs, it has a distinctive periplasmic β-jelly roll domain. BP951000_RS09575 exhibited the best structural alignment (Z-score = 8.6 and RMSD = 6 Å) with LptD (PDB ID: 5IXM) of *Yersinia pestis* (Table 2), validating the functional annotation of LptD in *B. pilosicoli.* Sequence variation analysis across nine strains of *B. pilosicoli* showed seven variations (N14, G137, I257, I382, E454, D600, G944) (Figure 3A, Tables-3, S3). When mapped onto the structural model, G944 was positioned in the ICL region, D600 in the ECL region, I382 and E454 in the TM region and the rest three variations in the β-jelly roll domain of the β-barrel structure (Table S3).

**Figure 3:**
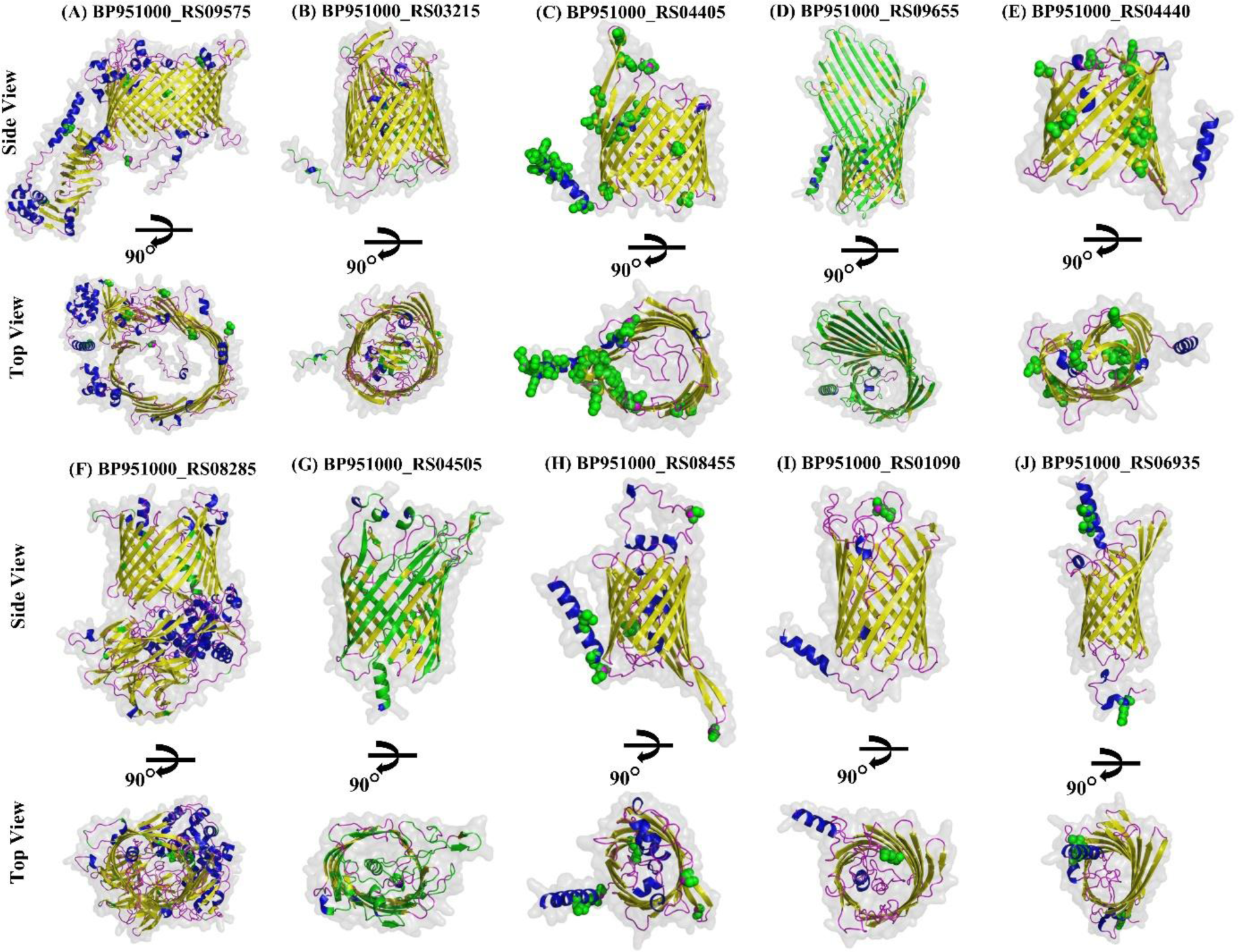
Structural models of β-barrel outer membrane proteins in Group B. Group B includes 29 proteins, of which 10 proteins are illustrated here. β-strands, α-helices and loops are shown in yellow, blue, and magenta colors, respectively. Green spheres indicate amino acid variations across nine strains of *B. pilosicoli*. Proteins exhibiting more than 40 variations are represented using ribbon representation in green colour.

#### 3.2.11 BP951000_RS03215

BP951000_RS03215 is annotated as TonB-dependent siderophore receptor in *B. pilosicoli*. SignalP predicted a secretory signal peptide, though LipoP predicted it as a cytoplasmic protein. The structural model revealed a β-barrel architecture comprising 22 β-strands at C-terminal and an N-terminal plug domain (Figure 3B). Structural homology search of BP951000_RS03215 using the DALI server revealed the best structural alignment with Vitamin B12 transporter BtuB (PDB ID: 3RGM) of *E. coli* K-12 (Z-score =32.1, RMSD = 2.7 Å) (Table 2). BtuB belongs to the TonB-dependent transporter (TBDT) family and consists of two domains: a 22-stranded TM β-barrel domain and an N-terminal globular luminal domain (also known as a hatch or plug domain) that occludes the lumen of the BtuB barrel.^106^ In *E. coli*, vitamin B12 (cobalamin) transport across the OM is mediated by the extracellular loops of BtuB.^106–108^ In addition to cobalamin, TBDTs also facilitate the transport of various molecules such as haem, ferric-siderophores, sucrose, and maltodextrin.^109,110^ Similarly, BP951000_RS03215 consists of both a 22-stranded β-barrel domain and an N-terminal plug domain. The highest Z-score of 32.1 indicates the true homology between BP951000_RS03215 and BtuB of *E. coli* K-12. Thus, BP951000_RS03215 might have a role in cobalamin transport across the OM. Sequence variation analysis across nine strains of *B. pilosicoli* identified 59 variations (Table S4) that have been mapped onto the structural model (Figure 3B).

#### 3.2.12 BP951000_RS04405

BP951000_RS04405 is annotated as Toxin A in the UniProt database.^20^ However, it is annotated as hypothetical protein in NCBI. This protein was predicted to have a secretory signal peptide with a cleavage position between residues 21 and 22. The structural model of BP951000_RS04405 revealed a β-barrel structure consisting of 18 β-strands, nine extracellular and eight periplasmic loops (Figure 3C). The protein showed the highest structural homology with probable porin (OccK7) (PDB ID: 4FRT) of *P. aeruginosa* PAO1(Z-score = 22.9 and RMSD = 3.5 Å) (Table 2). OccK protein family of *P. aeruginosa* shows the presence of a ladder of basic amino acids (Arg + Lys = 11 %) extending from the extracellular to the periplasmic surface, creating an energetically favorable environment for the small, carboxyl-containing substrates, facilitating their uptake into the cell to support its growth and function.^111–114^ BP951000_RS04405 showed a reduced arginine and lysine content (Arg + Lys = 7.6 %). It is plausible to suggest that BP951000_RS04405 could be a member of the OccK protein family and have a similar function, though it may not specifically transport negatively charged molecules. 23 amino acid sequence variations were identified across nine strains of *B. pilosicoli*: M1, H2, R3, I4, I6, L8, T9, M18, V19, T24, N32, S34, N41, F84, K90, N98, I101, S102, N104, S175, Q183, I264, T303 (Tables-3, S3).

When mapped onto the structural model, six variations (N41, F84, K90, N98, S175, Q183) in the ECL region, eight variations (T24, N32, S34, I101, S102, N104, I264, T303) in the TM β-barrel region and rest of the variations were found to be present in the periplasmic region of the protein (Figure 3C, Table S3).

#### 3.2.13 BP951000_RS09655

BP951000_RS09655, annotated as a hypothetical protein in NCBI, is annotated as DUF5723 domain-containing protein in the UniProt database. It carries a secretory signal peptide with a cleavage position between residues 18 and 19. The structural model revealed a 16-stranded β-barrel architecture in which β-strands 3 to 12 were longer than the rest of the β-strands, giving the barrel an asymmetrical shape on the extracellular side (Figure 3D). DALI search for structural homolog showed the best structural alignment with CymA protein (PDB ID: 4V3G) of *Klebsiella oxytoca* (Z-score= 17.7, RMSD = 6.5 Å) (Table 2). The 14-stranded CymA protein aligned structurally with only 14 β-strands of the 16-stranded β-barrel of BP951000_RS09655. CymA, an OM protein in Gram-negative bacteria, is a passive diffusion channel for large molecules (cyclodextrins and linear maltooligosaccharides).^115,116^ The pore diameter of BP951000_RS09655, comparable to CymA, suggests that it may function as a passive diffusion channel in *B. pilosicoli* OM for bulky molecule uptake. Structural alignment of BP951000_RS09655 with CymA using the US-Align server resulted in an RMSD value of 4.28 Å validating high structural homology. Amino acid sequence comparison of BP951000_RS09655 across nine strains of *B. pilosicoli* revealed 315 variations and many deletions, indicating that it is a highly variable protein (Table S4). These variations were spread throughout the protein (Figure 3D).

#### 3.2.14 BP951000_RS04440

BP951000_RS04440, a hypothetical protein, has a secretory signal peptide with a cleavage position between residues 17 and 18. The structural model generated by AlphaFold 3 revealed a 16-stranded β-barrel conformation with a lateral gate between strands 1 and 16 (Figure 3E). BP951000_RS04440 exhibited the best structural match with Translocation and Assembly Module protein TamA (PDB ID: 4C00) of *E. coli*, with a Z-score of 18.2 (RMSD = 4.2 Å) (Table 2). TamA is an Omp85 superfamily protein consisting of three N-terminal POTRA domains and a 16-stranded TM β-barrel region towards the C-terminal.^117^ TamA inserts autotransporter β-barrels into the membrane and translocates the passenger domain into the extracellular space.^117–119^ While the structural alignment between BP951000_RS04440 and TamA suggests a high degree of similarity in their TM β-barrel regions, a difference exists in the structures of both proteins. Unlike TamA, BP951000_RS04440 lacks POTRA domains. This indicates that BP951000_RS04440, despite its β-barrel resemblance, may not perform the specific functions associated with TamA. Thus, BP951000_RS04440 is predicted to be a structural homolog of TamA, with its functional role in *B. pilosicoli* requiring further characterization. Sequence variation across nine strains of *B. pilosicoli* revealed ten variations (K104, K113, S117, Y124, I132, T134, N151, G153, L243, V252) (Tables-3, S3). When mapped onto the structural model, two variations (K104 and L243) were present in the ECL region of the protein. In contrast, the rest of the variations were present in the TM β-barrel domain of the protein (Figure 3E, Table S3).

#### 3.2.15 BP951000_RS08285

BP951000_RS08285, annotated as a Trep protein in *B. pilosicoli*. Trep protein in *Pseudomonas fluorescens* catalyzes the phosphorylation of trehalose along with its translocation across the OM.^120^ LipoP predicted BP951000_RS08285 as a cytoplasmic protein, and SignalP predicted the absence of a signal peptide. The structural model generated by AlphaFold 3 revealed a C-terminal β-barrel structure consisting of 16 β-strands and a large N-terminal periplasmic domain (Figure 3F). It has a lateral opening between the 1^st^ and 16^th^ β-strands. Structure-based functional annotation of BP951000_RS08285 showed the best match (Z-score = 21 and RMSD = 2.5 Å) with TolB proteins of *E. coli* K-12 (3IAX) and *Citrobacter freundii* (PDB ID: 2IVZ) (Table 2). TolB of *E. coli* K-12 is a periplasmic protein with a distinct two-domain structure: an α/β N-terminal domain (residues 1–145), and a large C-terminal domain with a six-bladed β-propeller structure (Figure S4).^121^ It is involved in assembling outer membrane trimeric porins and maintains the integrity of the cell envelope.^122–124^ Structural similarity of the periplasmic domain of BP951000_RS08285 with C-terminal domain of TolB of *E. coli* K-12 suggests that it might perform similar functions. BP951000_RS08285 features an additional TM β-barrel domain absent in *E. coli* TolB. This domain suggests potential additional functions unique to *B. pilosicoli*. Upon using only the β-barrel domain of BP951000_RS08285 as a query for structural homology search resulted in filamentous hemagglutinin transporter protein FhaC (4QL0) of *Bordetella pertussis* as the top hit. FhaC consists of a C-terminal 16-stranded β-barrel and an N-terminal extension helix H1, connected by a 25-residue linker to two POTRA domains (Figure S4).^125^ In FhaC, the POTRA domains play a role in substrate recognition^126^, while the β-barrel serves as a translocation pore for the secretion of filamentous hemagglutinin (FHA)^127^, which contributes to virulence and biofilm formation in *B. pertussis*^128^. However, BP951000_RS08285 does not have POTRA domains. Structural similarity of BP951000_RS08285 to β-barrel domain of FhaC transporter protein suggests it might function as a translocation pore facilitating substrate transport. Furthermore, sequence homology search using BLASTp revealed that other spirochete genera (*Treponema*, *Borrelia*, and *Leptospira*) do not have a homolog of BP951000_RS08285. Multiple Sequence Alignment of BP951000_RS08285 homologs across nine strains of *B. pilosicoli* revealed 43 variations (Table S4).

#### 3.2.16 BP951000_RS04505

BP951000_RS04505 is annotated as a variable surface protein, VspH in *B. pilosicoli*.^20^ As previously described in section 3.2.3, Vsp proteins mediate adherence of the bacteria to the host cell.^60^ It has no signal peptide as predicted by SignalP while LipoP predicted the presence of TM helices at N-terminal. The structural model generated by AlphaFold 3 revealed a 16-stranded β-barrel architecture with eight extracellular loops and seven periplasmic loops (Figure 3G). Structural homology search showed the highest similarity to *E. coli* K-12 Poly-beta-1,6-N-acetyl-D-glucosamine export protein (PgaA, PDB ID: 4Y25) with a Z-score of 16.2 and an RMSD value of 3.2 Å (Table 2). As discussed in section 3.2.2, the role of PgaA implies that BP951000_RS04505 might have a role in translocation across the OM. *B. hyodysenteriae* VspH has been included in a potential vaccine.^65^ Sequence comparison across nine strains of *B. pilosicoli* revealed 247 variations and multiple deletions, indicating its high variability (Table S4). These variations were spread throughout the protein (Figure 3G).

#### 3.2.17 BP951000_RS08455

BP951000_RS08455 is classified as a PorV/PorQ family protein in the UniProt database. and as a hypothetical protein in NCBI. PorV and PorQ proteins are integral components of the Type IX secretion system (T9SS), the protein export mechanism of the Gram-negative Fibrobacteres-Chlorobi-Bacteroidetes superphylum. The T9SS system comprises over 15 proteins, with PorV as a β-barrel membrane protein.^129^ BP951000_RS08455 was predicted to have a lipoprotein signal peptide with a cleavage position between residues 21 and 22. The structural model revealed a 14-stranded β-barrel structure with seven extracellular loops (Figure 3H). BP951000_RS08455 exhibited best match (Z-score = 27.7 and RMSD = 3.5 Å) with long-chain fatty acid transport protein (FadL) (PDB ID: 2R88) of *E. coli* (Table 2). FadL has a monomeric 14-stranded β-barrel structure, with its interior blocked by an N-terminal hatch domain.^130^ Moreover, FadL has a distinct inward bend in one of its β-strands, forming a lateral opening in the TM barrel^130^, essential for transporting long-chain fatty acids.^131,132^ Although an N-terminal hatch domain also blocks the interior of BP951000_RS08455 (Figure 3H), but no lateral opening was observed in this protein, distinguishing it from FadL. Nonetheless, structural similarity to FadL suggests that BP951000_RS08455 might perform a comparable role, potentially in the transport of hydrophobic molecules across the membrane. PorV belongs to the FadL family of 14-stranded β-barrel OM porins. Similar to other FadL family members, the pore of PorV is obstructed by the N-terminal region of the protein. In previously characterized FadL family proteins, the third strand of the barrel kinks inwards to cover an N-terminal ‘hatch’ domain at the periplasmic end of the pore. However, in PorV, the N-terminal region occupies the pore, and the third and fourth β-strands bend outwards from the barrel axis. The inter-strand loop between these strands extends into the interior of the SprA barrel (protein-conducting channel of T9SS translocon) via a lateral opening.^133^ BP951000_RS08455 has an insertion at position 255 compared to its homologs found in nine strains included in this analysis. Sequence comparison across nine strains of *B. pilosicoli* showed six variations (L12, S20, N22, A117, R187, S253) (Figure 3H, Tables-3, S3). When mapped onto the structural model, two variations (N22 and A117) in ECL region, S253 variation in ICL region, R187 in the β-barrel TM region and remaining two variations were present in the N-terminal region of the protein (Table S3).

#### 3.2.18 BP951000_RS01090

BP951000_RS01090 is annotated as a hypothetical protein in NCBI; in contrast, the protein is annotated as a variable surface protein VspH in the UniProt database. It carries a secretory signal peptide with a cleavage position between residues 18 and 19. The structural model generated by AlphaFold 3 showed a 14 β-stranded β-barrel architecture (Figure 3I).

BP951000_RS01090 exhibited the best structural alignment (Z-score = 21, RMSD = 3.2 Å) with the CymA protein (PDB ID: 4V3G) of *K. oxytoca* (Table 2). Given the known function of CymA of *K. oxytoca* (discussed previously in section 3.2.13), it is plausible to speculate that BP951000_RS01090 might function as a diffusion channel for uptake of bulky molecules. Sequence alignment of BP951000_RS01090 across nine strains of *B. pilosicoli* showed variation at position E258, which was present in the ECL region (Figure 3I, Tables-3, S3). Structural alignment of BP951000_RS04760, BP951000_RS02055 and BP951000_RS04505 and BP951000_RS01090 using US-align server which resulted in an RMSD of 4 Å (Figure S5).

#### 3.2.19 BP951000_RS06935

BP951000_RS06935, is annotated as a hypothetical protein. It is predicted to carry a secretory signal peptide with a cleavage position between residues 21 and 22. The structural model of BP951000_RS06935 showed a 12-stranded β-barrel domain and six extracellular loops (Figure 3J). Structural homology search using BP951000_RS06935 model exhibited the best match with β-barrel domain of hemoglobin-binding protease hbp autotransporter (PDB ID: 3AEH) of *E. coli* (Table 2). It had a Z-score of 20.8 (RMSD = 3.2 Å). Hbp is a member of a sub-group of autotransporters called SPATEs (<u>s</u>erine <u>p</u>rotease <u>a</u>utotransporters of <u>E</u>nterobacteriaceae), consisting of a extracellular passenger domain at N-terminal in addition to a β-barrel domain at C-terminal.^134^ The β-barrel domain helps the passenger domain to secrete into the extracellular space, where it is released proteolytically from the cell. The passenger domain having serine protease activity degrades haemoglobin and helps uptake iron and heme from the host cell.^134–136^ BP951000_RS06935 lacks a periplasmic domain. However, homology between the β-barrels of BP951000_RS06935 and *E. coli* Hbp autotransporter suggests that it might be involved translocation of substances into the extracellular space. Sequence variation analysis across nine strains of *B. pilosicoli* showed four variations (S9, I10, V13, R298) (Figure 3J, Tables-3, S3). When mapped onto the structural model, these variations were present in the N-terminal and C-terminal region of the β-barrel structure (Tables S3).

#### 3.2.20 BP951000_RS11380

BP951000_RS11380, a hypothetical protein, contains a secretory signal peptide with a cleavage position between residues 19 and 20. However, it is annotated as ToxinA in the UniProt database. It has a TM helix at N-terminal. Structural model of BP951000_RS11380 revealed a β-barrel consisting of 12-anti-parallel β-strands, six extracellular and five periplasmic loops (Figure 4A). BP951000_RS11380 exhibited best structural alignment (Z-score = 17.6, RMSD = 2.8 Å) with Serine protease EspP (PDB ID: 2QOM) of *E. coli* O157:H7 (Table 2). EspP is a member of the <u>s</u>erine <u>p</u>rotease <u>a</u>utotransporters of <u>E</u>nterobacteriaceae (SPATE) family of autotransporters.^137^ Autotransporters are specialized porins produced by pathogenic Gram-negative bacteria, comprising a C-terminal β-barrel domain embedded in the membrane, creating a pore in the outer membrane. This pore allows the transport of an N-terminal passenger domain that functions as a virulence factor.

**Figure 4:**
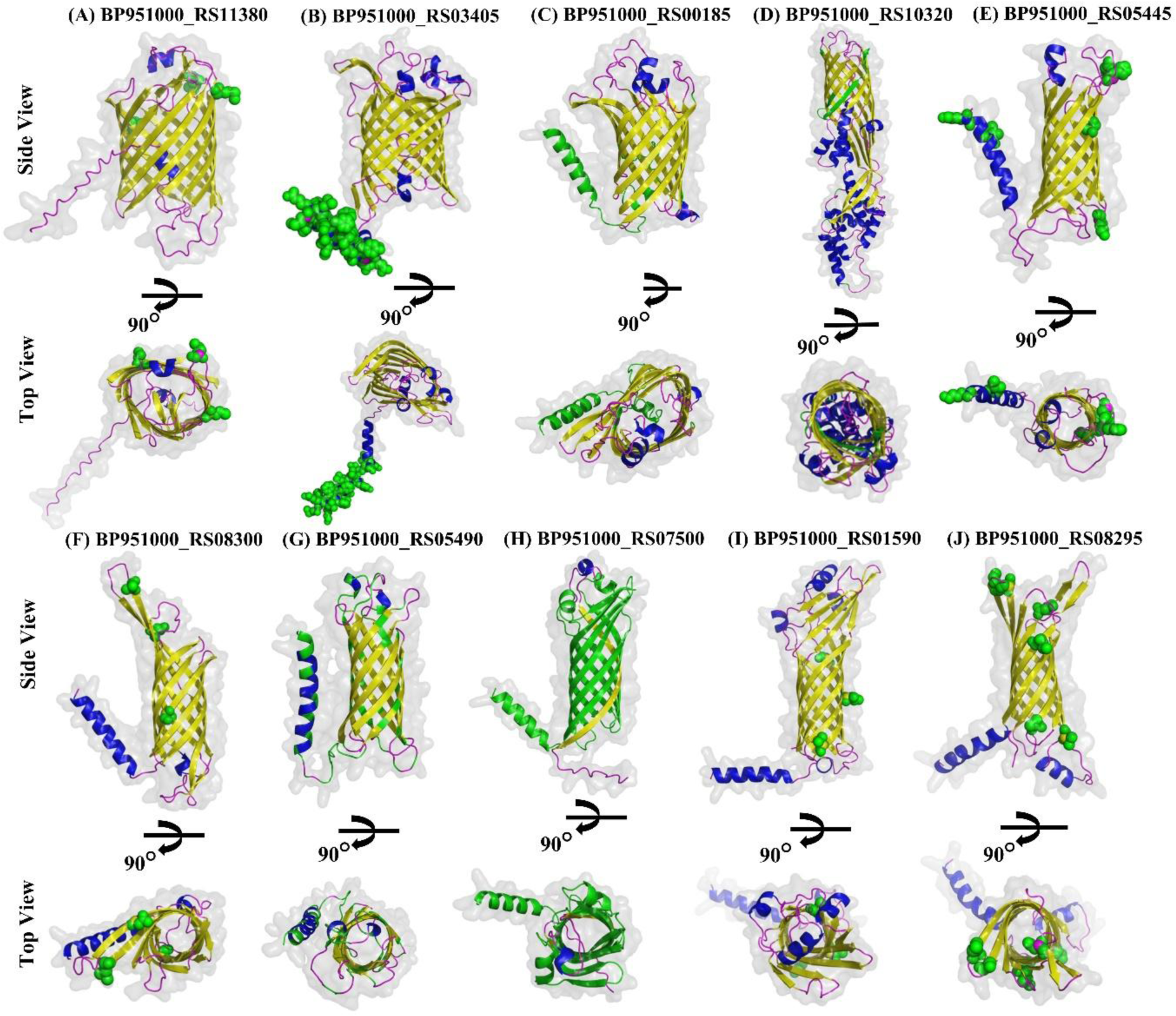
Structural models of β-barrel outer membrane proteins in Group B. Structural models of next 10 proteins of Group B are shown here, arranged in descending order by the number of β-strands in the β-barrel. β-strands, α-helices and loops are shown in yellow, blue, and magenta colors, respectively. Green spheres indicate amino acid variations across nine strains of *B. pilosicoli*. Proteins exhibiting more than 40 variations are represented using ribbon representation in green colour.

However, some autotransporters consist of only the autotransporter domain, with their upstream genes encoding the passenger or toxin protein.^138^ EspP passenger domain acts as serine protease that cleaves various mammalian proteins such as pepsin A and human coagulation factor V.^139–141^ In this context, the identification of BP951000_RS11380 as a structural homolog of EspP suggests that it might function as an autotransporter for passenger proteins with serine protease and virulence properties, which could be encoded either upstream of BP951000_RS11380 or elsewhere in the genome. Sequence comparison across nine strains of *B. pilosicoli* showed three variations (M126, M154, I278) (Figure 4A, Tables-3, S3). When mapped onto the structural model, M126 was present in the ECL region and other two variations (M154, I278) were found to be present in the TM β-barrel region of the protein (Table S3).

#### 3.2.21 BP951000_RS03405

BP951000_RS03405, annotated as a hypothetical protein, has a secretory signal peptide with a cleavage position between residues 19 and 20. AlphaFold 3-generated structural model revealed a 12-stranded β-barrel architecture, with six extracellular and five periplasmic loops (Figure 4B). A search using the DALI server revealed that BP951000_RS03405 exhibited the best structural alignment to Outer Membrane Phospholipase A (OMPLA) (PDB ID: 1ILD) from *E. coli*, with a Z-score of 18.1 (RMSD=2.7 Å) (Table 2). OMPLA is an acyl hydrolase that catalyzes the hydrolysis of acyl ester bonds present in phospholipids and lyso-phospholipids and is also involved in the secretion of colicins.^142–144^ Therefore, BP951000_RS03405, a structural homolog of *E. coli* OMPLA, may serve similar functions in *B. pilosicoli*. OMPLA has a unique His–Ser–Asn catalytic triad, responsible for its catalytic activity. Similar type of catalytic triad was not found in BP951000_RS03405, but Ser and Asn residues were present at positions 224 and 225, respectively. 24 amino acid sequence variations were identified across nine strains of *B. pilosicoli*: M1, R2, L3, K4, F5, F6, F7, L8, I9, F10, L11, F12, L13, S14, L15, S16, L17, Y18, T19, Q20, D21, N22, E23, A24 (Tables-3, S3). When mapped onto the structural model, all the variations were present in the periplasmic region of the β-barrel structure (Figure 4B, Tables S3).

#### 3.2.22 BP951000_RS00185

BP951000_RS00185, annotated as a hypothetical protein, carries a secretory signal peptide with a cleavage position between residues 20 and 21. The structural model of the protein revealed a 12-stranded β-barrel conformation consisting of six extracellular and five periplasmic loops (Figure 4C). BP951000_RS00185 showed the best structural match (Z-score = 18.3 and RMSD = 3.1 Å) with OMPLA (PDB ID: 1QD6) of *E. coli* K-12 (Table 2). As discussed in section 3.2.21, It catalyses the acyl ester bonds in phospholipids and lysophospholipids and the secretion of colicins.^142–144^ Therefore, BP951000_RS00185, a structural homolog of *E. coli* OMPLA, might have similar role in *B. pilosicoli*. OMPLA possesses a unique catalytic triad essential for its catalytic activity (discussed in section 3.2.21). In contrast, a similar catalytic triad was not identified in BP951000_RS03405; however, Ser-Asn residues were observed at positions 25-26 and 73-74. Sequence variation across nine strains of *B. pilosicoli* revealed 58 variations which have been mapped onto the structural model (Figure 4C, Table S4). These variations were mainly present in the signal peptide region of the protein (Figure 4C).

#### 3.2.23 BP951000_RS10320

BP951000_RS10320 is annotated as a hypothetical protein and possesses a secretory signal peptide. The structural model generated by AlphaFold 3 revealed a β-barrel configuration consisting of 10 anti-parallel β-strands, with several β-strands and α-helices extending towards the periplasmic side, creating an elliptical pore (Figure 4D). During structural homology search from the DALI server, the top four hits were synthetic constructs of a TM β-barrel, the fifth hit, i.e. OpcA (PDB ID: 2VDF) from *N. meningitidis* was considered as the best structural homolog (Z-score=14.7, RMSD =3 Å) (Table 2). OpcA is a 10-stranded β-barrel OMP that promotes the adhesion of the bacterium to the host epithelial surface through interaction with proteoglycan surface receptors.^145,146^ It is plausible to speculate that BP951000_RS10320, too, might function as an adhesin. 25 amino acid sequence variations were identified across nine strains of *B. pilosicoli*: L18, D48, E55, F251, G285, E400, Y401, G402, I403, F404, T405, K406, Q407, L408, A409, I410, S411, F412, I413, P414, I415, N416, I417, R418, F419 (Figure 4D, Tables-3, S3). Amongst these, no variation was present in the ECL region.

#### 3.2.24 BP951000_RS05445

BP951000_RS05445, annotated as DUF3575 domain-containing protein in the UniProt database carries a secretory signal peptide as predicted by SignalP. In contrast, it is annotated as a hypothetical protein in NCBI. LipoP, however, identified it as a cytoplasmic protein. AlphaFold 3-generated structural model revealed an eight-stranded β-barrel architecture (Figure 4E). BP951000_RS05445 exhibited the best structural homology with two proteins: OmpF (PDB ID: 4RLC) of *P. aeruginosa* PAO1 (Z-score = 16.1 and RMSD = 2.3 Å) and NspA (PDB ID: 1P4T) of *N. meningitidis* (Z-score = 16.1 and RMSD = 2.6 Å) (Table 2).

Given the known roles of OmpF and NspA in section 3.2.4 and 3.2.5 respectively, it is plausible to hypothesize that BP951000_RS05445 might serve similar roles in *B. pilosicoli*, potentially contributing to bacterial adhesion, immune system interaction, or even biofilm formation. Amino acid sequence variation across nine strains of *B. pilosicoli* revealed six variations (K2, I7, A79, N87, H89, K158) (Figure 4E, Tables-3, S3). When mapped onto the structural model, two variations (N87, H89) in the ECL region, K158 in the ICL region, A79 in the β-barrel domain and rest two were found to be present in the N-terminal region of the protein (Table S3).

#### 3.2.25 BP951000_RS08300

BP951000_RS08300, annotated as a tia invasion determinant, functions as both an adhesin and an invasin in enterotoxigenic *Escherichia coli* (ETEC) strains.^147^ Tia protein has eight TM amphipathic β-sheets with four loops that are exposed on the surface of the bacterial cell. Tia is an invasin and adhesin that binds a specific receptor on HCT8 human ileocecal epithelial cells.^148^ BP951000_RS08300 carries a secretory signal peptide, and its structural model reveals an elliptical eight-stranded β-barrel architecture, with two extended β-strands reaching towards the extracellular side (Figure 4F). BP951000_RS08300 exhibited the best structural alignment with NspA (PDB ID: 1P4T) from *N. meningitidis* with the highest Z-score of 17.9 (RMSD = 2 Å) (Table 2). Based on the role of NspA in *N. meningitidis* (described in section 3.2.5), BP951000_RS08300, another structural homolog, may perform similar functions. Sequence comparison across nine *B. pilosicoli* strains revealed three variations (L143, N156, S200) (Figure 4F, Tables-3, S3). When mapped onto the structural model, these variations were present in the β-barrel domain of the protein (Table S3).

#### 3.2.26 BP951000_RS05490

BP951000_RS05490 is annotated as a hypothetical protein in NCBI. However, it is annotated as tia invasion determinant in the UniProt database. It has a secretory signal peptide with a cleavage position between residues 23 and 24. LipoP, however, predicted it as a cytoplasmic protein. The structural model of BP951000_RS05490 revealed an eight-stranded β-barrel architecture (Figure 4G). BP951000_RS05490 showed the best structural match, based on the highest Z-score of 15.9 (RMSD = 2.5 Å), with NspA (PDB ID: 1P4T) of *N. meningitidis* (Table 2). BP951000_RS05490, another structural homolog of *N. meningitidis* NspA, may serve similar functions (discussed in section 3.2.5). Sequence comparison across nine strains of *B. pilosicoli* revealed 61 variations and deletions at two positions (Figure 4G, Table S4).

#### 3.2.27 BP951000_RS07500

BP951000_RS07500, annotated as a hypothetical protein, has a secretory signal peptide with a cleavage position between residues 19 and 20. The structural model of BP951000_RS07500 showed an eight-stranded β-barrel structure and four extracellular loops (Figure 4H).

BP951000_RS07500 showed the best structural match with the β-barrel domain of *E. coli* K-12 OmpA (PDB ID: 9FZC) (Z-score =15.1, RMSD = 3Å) (Table 2). However, no periplasmic domain similar to that of OmpA (discussed in section 3.2.9) is present in BP951000_RS0750. As described previously in section 3.2.9, OmpA of *E. coli* K-12 serves multiple functions, acting as a receptor for bacteriophages, facilitating the action of colicins vesicles, mediating F-dependent conjugation and maintaining the structural integrity of the membrane. It also acts as a diffusion channel for small-sized solutes and mediates the virulence and pathogenicity of *E. coli*.^96–104^ It is plausible to speculate that BP951000_RS07500, a structural homolog of the *E. coli* OmpA β-barrel domain, may perform similar functions. Sequence comparison of BP951000_RS07500 across nine strains of *B. pilosicoli* revealed that it is a highly variable protein with 183 variations across the polypeptide chain length of 232 residues (Figure 4H, Table S4). These variations were localized in 1^st^ to 7^th^ strand of the TM β-barrel protein (Figure 4H).

#### 3.2.28 BP951000_RS01590

BP951000_RS01590, annotated as a hypothetical protein, contains a secretory signal peptide with a cleavage site between residues 24 and 25. Additionally, LipoP has identified a TM helix at the N-terminal of the protein. The structural model of BP951000_RS01590 revealed an eight-stranded β-barrel architecture with four extracellular loops (Figure 4I). The loops extending into the extracellular region comprises five short β-strands and three short α-helices. BP951000_RS01590 exhibited the best match (Z-score = 13.7 and RMSD = 3 Å) with the outer membrane protein OprG (PDB ID: 2X27) of *P. aeruginosa* (Table 2). OprG is an eight-stranded OM β-barrel protein belonging to the OmpW family.^149^ The OmpW family of small outer membrane proteins is commonly found in Gram-negative bacteria and is involved in the breakdown of small, hydrophobic compounds like medium-chain alkanes (AlkL) and naphthalene (NahQ and DoxH).^150–152^ Additionally, OmpW could mediate diffusion of hydrophobic molecules.^153^ This suggests that BP951000_RS01590 might have a role in the utilization and uptake of hydrocarbons and mediate the transport of hydrophobic molecules. Sequence variation across nine strains of *B. pilosicoli* revealed three variations (V205, I215, V221) (Figure 4I, Tables-3, S3). When mapped onto the structural model, these variations were present in the TM barrel domain of the protein (Figure 4I).

#### 3.2.29 BP951000_RS08295

BP951000_RS08295, another tia invasion determinant, functions as an adhesin and an invasin (described in section 3.2.25).^147^ It possesses a secretory signal peptide. The structural model of BP951000_RS08295 revealed an eight-stranded β-barrel structure of which two β-strands were longer, forming an elliptical pore-like structure on the extracellular side (Figure 4J). BP951000_RS08295 exhibited the best structural alignment, based on the highest Z-score of 19.4 (RMSD = 2.3 Å), with NspA (PDB ID: 1P4T) of *N. meningitidis* (Table 2). In this light, our discovery of BP951000_RS08295 as an NspA structural homolog indicates it might perform similar functions like NspA. Sequence comparison across nine strains of *B. pilosicoli* revealed six variations (N34, I49, V123, S141, I144, V167) (Figure 4J, Tables-3, S3). When mapped onto the structural model, N34 was present in the ECL region, while the rest of the variations were in the β-barrel domain of the protein (Table S3).

#### 3.2.30 BP951000_RS08975

BP951000_RS08975 has been annotated in the NCBI database as a TonB-dependent receptor domain-containing protein (TBDRs).^20^ However, it is annotated as Ser/Thr protein kinase in the UniProt database. TBDRs facilitate the transport of specific substrates across the outer membrane by utilizing energy from the proton motive force, which is transmitted by the TonB−ExbB−ExbD complex located in the inner membrane.^109,154^ SignalP predicted a secretory signal peptide in this protein. The AlphaFold 3-generated structural model revealed a 13-stranded incomplete β-barrel structure, forming approximately half of a β-barrel structure (Figure 5A). It is possible that this incomplete 13-stranded β-barrel could form a dimer or a higher-order oligomer. However, an attempt to predict a dimeric structure using the AlphaFold 3 server was inconclusive, with pTM and ipTM scores of 0.35 and 0.19, respectively, which were too low to suggest a stable oligomeric assembly confidently. We also generated structural models for BP951000_RS08975 using other modelling tools (ESMFold, SWISS-MODEL, RoseTTA and TrRosetta) to validate the structure generated by AlphaFold 3. Structural models were aligned well with AlphaFold 3 prediction (RMSD = 4.61 Å). Structure-based functional annotation study of BP951000_RS08975 using the DALI server generated its best match with Vitamin B12 transporter BtuB (PDB ID: 2GSK) of *E. coli* (Z-score =18.2, RMSD =7.2 Å) (Table 2). As described in section 3.2.11, BtuB consists of two domains: a 22-stranded TM β-barrel domain and an N-terminal globular luminal domain (also known as a hatch or plug domain) that occludes the lumen of the BtuB barrel.^106^

**Figure 5:**
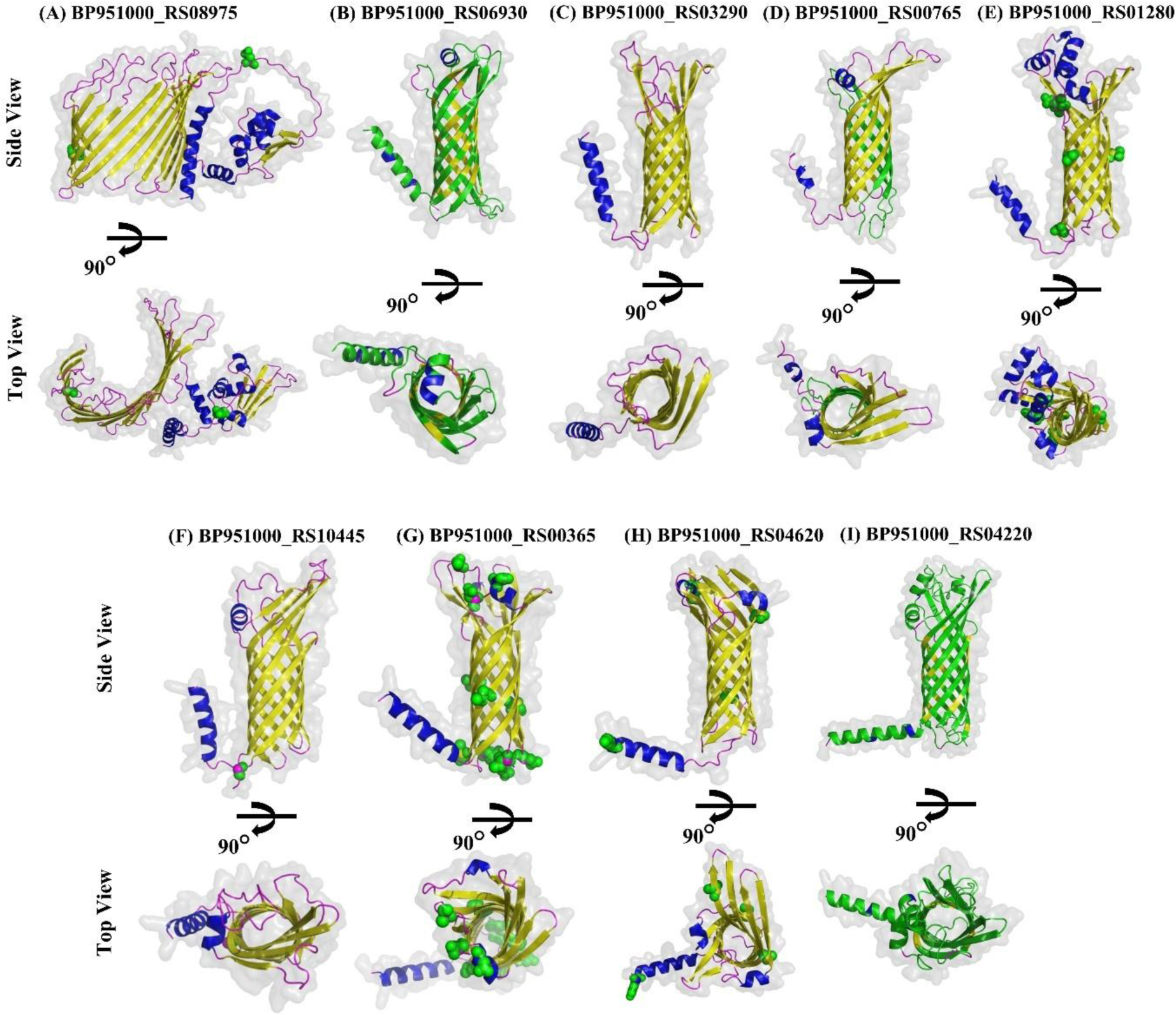
Structural models of β-barrel outer membrane proteins in Group B. Structural models of remaining 9 nine proteins of Group B are shown here. Proteins are arranged according to decreasing number of β-strands. Structural model of BP951000_RS08975 revealed an incomplete β-barrel structure consisting of 13 β-strands. β-strands, α-helices and loops are shown in yellow, blue, and magenta colors, respectively. Green spheres indicate amino acid variations across nine strains of *B. pilosicoli*. Proteins exhibiting more than 40 variations are represented using ribbon representation in green colour.

Despite a notable Z-score of 18.2, high RMSD value of 7.2 Å highlighted significant structural differences between BP951000_RS08975 and BtuB, suggesting that BP951000_RS08975 is unlikely to be a structural homolog of BtuB. Furthermore, sequence homology search using BLASTp revealed that BP951000_RS08975 does not have a homolog in other spirochete genera (*Treponema*, *Borrelia*, and *Leptospira*). Sequence comparison of BP951000_RS08975 across nine strains of *B. pilosicoli* revealed amino acid variations at two positions (D32, T371), of which one variation (D32) was found to be present in the ECL region (Figure 5A). The remaining one (T371) was present in the β-barrel TM region of the protein (Figure 5A, Tables-3, S3).

#### 3.2.31 Eight-stranded β-barrel proteins annotated as Serpentine receptor domain containing proteins

BP951000_RS01280, BP951000_RS04620, BP951000_RS00765, BP951000_RS03290, BP951000_RS04220, BP951000_RS10445, BP951000_RS00365 and BP951000_RS06930 are annotated as Serpentine receptor domain containing proteins in the UniProt database; in contrast, these are annotated as hypothetical proteins in NCBI. These proteins were predicted to have a signal peptide. Five Serpentine receptor domain containing proteins categorized under Group A have been discussed in section 3.2.9. AlphaFold 3-generated structural models for these proteins showed a β-barrel architecture comprising eight anti-parallel β-strands, four extracellular and three periplasmic loops (Figure 5B-5I). BP951000_RS01280, BP951000_RS00365 and BP951000_RS06930 showed the best structural alignment with NspA (PDB ID: 1P4T) of *N. meningitidis* (Table 2), with the highest Z-scores (ranging from 14 to 17). BP951000_RS04620, BP951000_RS00765, BP951000_RS03290, BP951000_RS04220 and BP951000_RS10445 revealed the best structural match with N-terminal β-barrel domain of OmpA (PDB ID: 9FZC) of *E. coli* K-12, in terms of highest Z-score (ranging from 14 to 17). As discussed in section 3.2.5, NspA is an eight-stranded β-barrel protein which is involved in attachment of the bacteria to the host immune system and promotes their colonization. It is a potential vaccine candidate in *N. meningitidis*.^82–85^ As described in section 3.2.9, *E. coli* OmpA consists of an N-terminal eight-stranded β-barrel domain and a periplasmic C-terminal domain. It functions as a receptor for bacteriophages, facilitates colicin vesicles action, mediates F-dependent conjugation, maintains membrane integrity, and serves as a diffusion channel for small solutes.^96–103^ It also contributes to virulence and pathogenicity of *E. coli*, making it a key target in the immune response.^104^ Given the known roles of OmpA and NspA, these proteins might perform analogous roles in *B. pilosicoli*. However, experimental evidence would be required for the same. Sequence comparison across nine strains of *B. pilosicoli* revealed variations at several positions which are listed in Table 3, S3 and S4.

## 4. Conclusion

The prevalence of HIS in developed nations ranges from 1.1% to 5%^155^, with a significantly higher rate (54%) among homosexual men and HIV-positive individuals^156^. Clinical diagnosis still relies on the histopathological assessment of biopsy samples^8^, and no human or animal vaccines are available. Hence, there is an urgent need to develop targeted diagnostic tools and effective therapeutic and preventive strategies. *B. pilosicoli* colonizes by attaching itself to the surface epithelium of the gastrointestinal (GI) tract of the host and causes IS in domesticated animals and humans. However, a limited understanding of the disease and its diverse clinical presentation underscore the need for focused research to elucidate its pathogenesis. *B. pilosicoli* is a diderm bacteria. Consequently, its outer membrane must play pivotal roles in selective permeability, nutrient uptake, adhesion, host-pathogen interactions, disease establishment, and immune evasion. However, there remains a significant gap in our knowledge of the *B. pilosicoli* OMP repertoire, collectively known as the OMPome, and its role in IS pathogenesis. In this study, we predicted and characterized 42 outer membrane proteins (OMPs) with TM β-barrel structures using a computational framework and consensus-based algorithm. Among these, 35 were previously annotated in the NCBI or UniProt databases, while seven were unannotated hypothetical proteins, none of which have been experimentally characterized. Structural modelling using AlphaFold 3 validated the β-barrel predictions. Structural homology between the predicted OMPs and known OMPs from other spirochetes and model Gram-negative bacteria led to functional inferences. Protein sequence variation analysis across nine *B. pilosicoli* strains revealed 12 proteins to be highly variable, containing more than 40 variations. High sequence variability could indicate dynamism in immune evasion mechanisms and high adaptability to the host environment. These findings have expanded our knowledge of the *B. pilosicoli* OMP repertoire, offering promising avenues for developing diagnostic, prophylactic and therapeutic strategies.

## Acknowledgements

AP is a recipient of junior research fellowship from the Department of Biotechnology, Government of India.

## Author contributions statement

**Amisha Panda:** methodology, software, investigation, data analysis, data curation, writing— original draft preparation. **Jahnvi Kapoor:** data analysis, visualization, writing—reviewing and editing. **B. Hareramadas**: visualization, writing—reviewing and editing. **Ilmas Naqvi**: visualization, writing—reviewing and editing. **Ravindresh Chhabra**: data analysis, writing—original draft preparation, writing—reviewing and editing. **Sanjiv Kumar:** conceptualization, methodology, software, writing—reviewing and editing, supervision.

**Anannya Bandyopadhyay**: methodology, data analysis, writing—original draft preparation, writing—reviewing and editing, supervision. All authors reviewed the final draft of the manuscript. All authors contributed to the manuscript revision, read and approved the submitted version.

## Data Availability Statement

The data supporting this study’s findings are available from the corresponding author upon reasonable request.

