## Supplemental material for "Identification and Characterization of Novel Outer Membrane Proteins of *Brachyspira pilosicoli*"

**Supplementary Table 1**: Comprehensive results from computational tools employed in our study to predict outer membrane β-barrel proteins.

| **Protein Accession No.** | **Locus Identifier** | **Protein name^#^** | **Pepstats** | | **SignalP** | **Cleavage Position** | **LipoP** | **DEG Match** | **PSORTb** | | **CELLO** | **OMPdb Match** | **MCMBB** | **TMBETADISC.RBF_AADP** | **Tmbed** | |
| --- | --- | --- | --- | --- | --- | --- | --- | --- | --- | --- | --- | --- | --- | --- | --- | --- |
|  |  |  | **Length** | **Mol.Wt. (kDa)** |  |  |  |  | **Localization** | **Score** |  |  |  |  | **Tmbed Count_B** | **Tmbed Count_b** |
| **GROUP A** |  |  |  |  |  |  |  |  |  |  |  |  |  |  |  |  |
| WP_013244106.1 | BP951000_RS05730 | Outer membrane protein assembly factor BamA | 872 | 100.1 | SP | CS pos: 24-25 | SpI | DEG10280066 | Outer Membrane | 9.92 | Outer Membrane | 500 | 0.005 | Outer Membrane Protein | 73 | 77 |
| WP_013244995.1 | BP951000_RS10215 | Variable surface protein, VspE | 367 | 40.0 | SP | CS pos: 21-22 | SpI | - | Unknown | 2 | Outer Membrane | 163 | 0.046 | Outer Membrane Protein | 70 | 82 |
| WP_013243917.1 | BP951000_RS04760 | Variable surface protein, VspD | 392 | 42.8 | SP | CS pos: 21-22 | SpI | - | Unknown | 2 | Outer Membrane | 205 | 0.039 | Outer Membrane Protein | 56 | 69 |
| WP_013243193.1 | BP951000_RS01125 | CsgG/HfaB family protein | 489 | 55.1 | LIPO | CS pos: 14-15 | CYT | - | Unknown | 5.48 | Outer Membrane | 13 | 0.018 | Outer Membrane Protein | 40 | 40 |
| WP_013243655.1 | BP951000_RS03440 | Outer membrane beta-barrel protein | 240 | 26.4 | SP | CS pos: 21-22 | SpI | - | Unknown | 4.72 | Outer Membrane | 5 | 0.058 | Outer Membrane Protein | 39 | 42 |
| WP_013244081.1 | BP951000_RS05600 | TolC family protein | 500 | 57.5 | SP | CS pos: 26-27 | SpI | - | Cytoplasmic | 8.96 | Outer Membrane | 52 | 0.002 | Outer Membrane Protein | 31 | 28 |
| WP_013244750.1 | BP951000_RS09000 | TolC family protein | 484 | 54.7 | SP | CS pos: 20-21 | SpI | - | Unknown | 2.5 | Outer Membrane | 54 | 0.035 | Outer Membrane Protein | 30 | 28 |
| WP_041747714.1 | BP951000_RS06235 | TolC family protein | 452 | 51.6 | SP | CS pos: 21-22 | SpI | - | Outer Membrane | 10 | Outer Membrane | 58 | 0.048 | Outer Membrane Protein | 20 | 20 |
| WP_013243940.1 | BP951000_RS04880 | Serpentine receptor domain containing protein | 214 | 23.4 | SP | CS pos: 21-22 | SpI | - | Outer Membrane | 9.52 | Outer Membrane | 5 | 0.03 | Outer Membrane Protein | 39 | 42 |
| WP_013243377.1 | BP951000_RS02055 | Serpentine receptor domain containing protein | 256 | 29.3 | SP | CS pos: 20-21 | SpI | - | Outer Membrane | 9.52 | Outer Membrane | 5 | 0.045 | Outer Membrane Protein | 42 | 44 |
| WP_013243376.1 | BP951000_RS02050 | Serpentine receptor domain containing protein | 254 | 28.6 | SP | CS pos: 20-21 | SpI | - | Outer Membrane | 9.49 | Outer Membrane | 1 | 0.043 | Outer Membrane Protein | 40 | 43 |
| WP_013244459.1 | BP951000_RS07540 | Serpentine receptor domain containing protein | 213 | 23.4 | SP | CS pos: 21-22 | SpI | - | Outer Membrane | 9.52 | Outer Membrane | 5 | 0.031 | Outer Membrane Protein | 40 | 37 |
| WP_013242998.1 | BP951000_RS00180 | Serpentine receptor domain containing protein | 224 | 25.0 | SP | CS pos: 20-21 | SpI | - | Unknown | 2 | Outer Membrane | 5 | 0.065 | Outer Membrane Protein | 40 | 42 |
| **GROUP B­­­** |  |  |  |  |  |  |  |  |  |  |  |  |  |  |  |  |
| WP_041747843.1 | BP951000_RS09575 | LPS-assembly protein LptD | 970 | 111.9 | SP | CS pos: 19-20 | SpI | DEG10200216 | Outer Membrane | 9.52 | Outer Membrane | 0 | 0.036 | Outer Membrane Protein | 111 | 118 |
| WP_041747581.1 | BP951000_RS03215 | TonB-dependent siderophore receptor | 649 | 75.2 | SP | CS pos: 18-19 | CYT | - | Outer Membrane | 9.49 | Outer Membrane | 1 | 0.015 | Non-Outer Membrane Protein | 91 | 102 |
| WP_013243854.1 | BP951000_RS04405 | ToxinA | 354 | 42.3 | SP | CS pos: 21-22 | SpI | - | Outer Membrane | 9.52 | Outer Membrane | 0 | 0.029 | Outer Membrane Protein | 68 | 76 |
| WP_013244879.1 | BP951000_RS09655 | DUF5723 domain-containing protein | 494 | 53.8 | SP | CS pos: 20-21 | SpI | - | Unknown | 2.5 | Outer Membrane | 0 | 0.038 | Outer Membrane Protein | 73 | 73 |
| WP_013243861.1 | BP951000_RS04440 | Hypothetical protein | 334 | 38.5 | SP | CS pos: 17-18 | SpI | - | Unknown | 4.72 | Outer Membrane | 0 | 0.018 | Outer Membrane Protein | 60 | 63 |
| WP_013244607.1 | BP951000_RS08285 | Trep protein | 926 | 108.1 | OTHER | None | CYT | - | Unknown | 4.69 | Outer Membrane | 0 | 0.021 | Outer Membrane Protein | 70 | 75 |
| WP_015274839.1 | BP951000_RS04505 | Variable surface protein VspH | 423 | 46.8 | OTHER | None | TMH | - | Unknown | 2 | Outer Membrane | 205 | 0.024 | Non-Outer Membrane Protein | 68 | 71 |
| WP_013244641.1 | BP951000_RS08455 | PorV/PorQ family protein | 334 | 36.8 | LIPO | CS pos: 21-22 | TMH | - | Unknown | 4.72 | Outer Membrane | 0 | 0.049 | Outer Membrane Protein | 49 | 56 |
| WP_013243185.1 | BP951000_RS01090 | Variable surface protein, VspH | 361 | 40.6 | SP | CS pos: 18-19 | SpI | - | Outer Membrane | 9.52 | Outer Membrane | 0 | 0.023 | Outer Membrane Protein | 64 | 68 |
| WP_013244339.1 | BP951000_RS06935 | hypothetical protein | 302 | 35.1 | SP | CS pos: 21-22 | SpI | - | Cytoplasmic Membrane | 9.82 | Outer Membrane | 0 | 0.016 | Outer Membrane Protein | 52 | 56 |
| WP_041747935.1 | BP951000_RS11380 | ToxinA | 289 | 33.1 | SP | CS pos: 19-20 | TMH | - | Unknown | 2 | Outer Membrane | 0 | 0.025 | Outer Membrane Protein | 46 | 59 |
| WP_013243647.1 | BP951000_RS03405 | Hypothetical protein | 327 | 39.3 | SP | CS pos: 19-20 | SpI | - | Unknown | 2 | Outer Membrane | 0 | 0.006 | Outer Membrane Protein | 51 | 54 |
| WP_013242999.1 | BP951000_RS00185 | Hypothetical protein | 300 | 34.4 | SP | CS pos: 20-21 | SpI | - | Outer Membrane | 9.49 | Outer Membrane | 0 | 0.031 | Outer Membrane Protein | 57 | 58 |
| WP_041747873.1 | BP951000_RS10320 | Hypothetical protein | 419 | 46.7 | SP | CS pos: 20-21 | SpI | - | Unknown | 2 | Outer Membrane | 0 | 0.023 | Outer Membrane Protein | 48 | 45 |
| WP_013244050.1 | BP951000_RS05445 | DUF3575 domain-containing protein | 201 | 22.5 | SP | CS pos: 21-22 | CYT | - | Unknown | 2 | Outer Membrane | 0 | 0.025 | Outer Membrane Protein | 34 | 36 |
| WP_013244610.1 | BP951000_RS08300 | Tia invasion determinant | 205 | 23.5 | SP | CS pos: 21-22 | SpI | - | Unknown | 2 | Outer Membrane | 12 | 0.025 | Non-Outer Membrane Protein | 33 | 37 |
| WP_013244059.1 | BP951000_RS05490 | Tia invasion determinant | 202 | 22.9 | SP | CS pos: 23-24 | CYT | - | Unknown | 2 | Outer Membrane | 0 | 0.023 | Outer Membrane Protein | 33 | 38 |
| WP_187287137.1 | BP951000_RS07500 | Hypothetical protein | 232 | 25.5 | SP | CS pos: 19-20 | SpI | - | Cytoplasmic Membrane | 9.82 | Outer Membrane | 0 | 0.032 | Outer Membrane Protein | 37 | 42 |
| WP_181893515.1 | BP951000_RS01590 | Hypothetical protein | 270 | 29.4 | SP | None | TMH | - | Cytoplasmic Membrane | 9.82 | Outer Membrane | 0 | 0.036 | Outer Membrane Protein | 41 | 41 |
| WP_228369485.1 | BP951000_RS08295 | Tia invasion determinant | 198 | 22.9 | SP | CS pos: 18-19 | SpI | - | Unknown | 4.69 | Inner Membrane | 8 | 0.013 | Non-Outer Membrane Protein | 34 | 37 |
| WP_013244745.1 | BP951000_RS08975 | TonB-dependent receptor domain-containing protein | 445 | 51.5 | SP | CS pos: 16-17 | CYT | - | Unknown | 5.48 | Outer Membrane | 0 | 0.031 | Outer Membrane Protein | 53 | 63 |
| WP_013244338.1 | BP951000_RS06930 | Serpentine receptor domain containing protein | 211 | 24.3 | OTHER | None | SpI | - | Cytoplasmic Membrane | 9.82 | Inner Membrane | 0 | 0.03 | Outer Membrane Protein | 36 | 40 |
| WP_014936494.1 | BP951000_RS03290 | Serpentine receptor domain containing protein | 222 | 25.6 | SP | CS pos: 23-24 | SpI | - | Cytoplasmic Membrane | 9.86 | Outer Membrane | 0 | 0.007 | Outer Membrane Protein | 40 | 41 |
| WP_014933009.1 | BP951000_RS00765 | Serpentine receptor domain containing protein | 232 | 26.4 | SP | CS pos: 21-22 | SpI | - | Outer Membrane | 9.49 | Outer Membrane | 0 | 0.046 | Outer Membrane Protein | 40 | 41 |
| WP_013243225.1 | BP951000_RS01280 | Serpentine receptor domain containing protein | 256 | 28.4 | SP | CS pos: 20-21 | SpI | - | Outer Membrane | 9.52 | Outer Membrane | 0 | 0.052 | Outer Membrane Protein | 40 | 42 |
| WP_013245039.1 | BP951000_RS10445 | Serpentine receptor domain containing protein | 230 | 25.6 | SP | CS pos: 21-22 | TMH | - | Outer Membrane | 9.49 | Outer Membrane | 0 | 0.04 | Outer Membrane Protein | 39 | 39 |
| WP_013243037.1 | BP951000_RS00365 | Serpentine receptor domain containing protein | 226 | 25.5 | SP | CS pos: 18-19 | SpI | - | Unknown | 2.5 | Outer Membrane | 0 | 0.061 | Outer Membrane Protein | 37 | 41 |
| WP_013243896.1 | BP951000_RS04620 | Serpentine receptor domain containing protein | 234 | 25.7 | SP | CS pos: 23-24 | SpI | - | Outer Membrane | 9.52 | Outer Membrane | 0 | 0.04 | Outer Membrane Protein | 40 | 42 |
| WP_013243815.1 | BP951000_RS04220 | Serpentine receptor domain containing protein | 254 | 28.6 | SP | CS pos: 23-24 | TMH | - | Cytoplasmic Membrane | 9.82 | Outer Membrane | 0 | 0.043 | Outer Membrane Protein | 39 | 39 |

#Proteins were named as per their annotation in NCBI and UniProt databases, searched using Protein Accession Number.

**Supplementary Table 2:** *B. pilosicoli* nine strains searched for predicted OMP’s variations compared to the reference genome 95/1000.

| Organism name | Strain | Assembly | Level | Size (Mb) | GC% | Scaffolds | CDS^#^ |
| --- | --- | --- | --- | --- | --- | --- | --- |
| *B. pilosicoli* | P43/6/78 | GCA_000325665.1 | Complete | 2.556 | 28 | 1 | 2208 |
| *B. pilosicoli* | MEI7141 | GCA_030168385.1 | Complete | 2.596 | 28 | 1 | 2295 |
| *B. pilosicoli* | MEI4046 | GCA_030168405.1 | Complete | 2.595 | 28 | 1 | 2290 |
| *B. pilosicoli* | MEI4028 | GCA_030168425.1 | Complete | 2.592 | 28 | 1 | 2287 |
| *B. pilosicoli* | ZH1243 | GCA_030168345.1 | Complete | 2.578 | 28 | 1 | 2261 |
| *B. pilosicoli* | ZH1145 | GCA_030168365.1 | Complete | 2.551 | 28 | 1 | 2226 |
| *B. pilosicoli* | ZH1268 | GCA_030168305.1 | Complete | 2.539 | 28 | 1 | 2209 |
| *B. pilosicoli* | ZH1244 | GCA_030168325.1 | Complete | 2.528 | 28 | 1 | 2220 |
| *B. pilosicoli* | B2904 | GCA_000296575.1 | Complete | 2.765 | 28 | 1 | 2658 |

^#^CDS: Coding Sequences

**Supplementary Table 3:** Sequence variations among nine strains in the Intracellular loops, extracellular loops, β-barrel TM region, and other regions of predicted OM β-barrel proteins.

| **Locus Identifier** | **Protein name^#^** | **Total variations** | **Variations in ICL region** | **Variations in ECL region** | **Variations in TM region** | **Variations in**  **other region** |
| --- | --- | --- | --- | --- | --- | --- |
| **GROUP A** |  |  |  |  |  |  |
| BP951000_RS05730 | Outer membrane protein assembly factor BamA | D60, A184, V465, A467, F512 | None | None | V465, A467, F512 | D60, V184 |
| BP951000_RS10215^η^ | Variable surface protein, VspE | 208 variations | Not determined^µ^ | Not determined^µ^ | Not determined^µ^ | Not determined^µ^ |
| BP951000_RS04760^η^ | Variable surface protein, VspD | 260 variations | Not determined^µ^ | Not determined^µ^ | Not determined^µ^ | Not determined^µ^ |
| BP951000_RS01125 | CsgG/HfaB family protein | S63, D79, T190, I210, L380 | None | None | L380 | S63, D79, T190, I210 |
| BP951000_RS03440 | Outer membrane beta-barrel protein | F24, V47, V64, N110, D169, A197 | None | V47 | F24, V64, N110, A197 | D169 |
| BP951000_RS05600 | TolC family protein | T246, N499 | None | None | None | T246, N499 |
| BP951000_RS09000 | TolC family protein | S90, S131 | None | S90 | S131 | None |
| BP951000_RS06235 | TolC family protein | K2, N3, F5, V6, F7, I8, I10, L12, S16, S25, N33, I42, E43, L93, S105, E136, I137, T210 | None | L93 | S105 | K2, N3, F5, V6, F7, I8, I10, L12, S16, S25, N33, I42, E43, E136, I137, L210 |
| BP951000_RS04880 | Serpentine receptor domain containing protein | N69 | N69 | None | None | None |
| BP951000_RS02055 | Serpentine receptor domain containing protein | H101, N163, M235 | None | N163, M235 | H101 | None |
| BP951000_RS02050 | Serpentine receptor domain containing protein | A27, L28, T108, A228, I247 | None | T108, A228 | A27, L28, I247 | None |
| BP951000_RS07540 | Serpentine receptor domain containing protein | M1, K2, K3, I4, I5, L6 | None | None | None | M1, K2, K3, I4, I5, L6 |
| BP951000_RS00180 | Serpentine receptor domain containing protein | I64, M115 | None | None | I64, M115 | None |
| **GROUP B** |  |  |  |  |  |  |
| BP951000_RS09575 | LPS-assembly protein LptD | N14, G137, I257, I382, E454, D600, G944 | G944 | D600 | I382, E454 | N14, G137, I257 |
| BP951000_RS03215^η^ | TonB-dependent siderophore receptor | 59 variations | Not determined^µ^ | Not determined^µ^ | Not determined^µ^ | Not determined^µ^ |
| BP951000_RS04405 | ToxinA | M1, H2, R3, I4, I6, L8, T9, M18, V19, T24, N32, S34, N41, F84, K90, N98, I101, S102, N104, S175, Q183, I264, T303 | None | N41, F84, K90, N98, S175, Q183 | T24, N32, S34, I101, S102, N104, I264, T303 | M1, H2, R3, I4, I6, L8, T9, M18, V19, |
| BP951000_RS09655^η^ | DUF5723 domain-containing protein | 315 variations | Not determined^µ^ | Not determined^µ^ | Not determined^µ^ | Not determined^µ^ |
| BP951000_RS04440 | Hypothetical protein | K104, K113, S117, Y124, I132, T134, N151, G153, L243, V252, L254, S308, N321 | None | K104, L243 | K113, S117, Y124, I132, T134, N151, G153, V252, L254, N321 S308, | None |
| BP951000_RS08285^η^ | Trep protein | 43 variations | Not determined^µ^ | Not determined^µ^ | Not determined^µ^ | Not determined^µ^ |
| BP951000_RS04505^η^ | Variable surface protein VspH | 247 variations | Not determined^µ^ | Not determined^µ^ | Not determined^µ^ | Not determined^µ^ |
| BP951000_RS08455 | PorV/PorQ family protein | L12, S20, N22, A117, R187, S253 | S253 | N22, A117, | R187 | L12, S20, |
| BP951000_RS01090 | Variable surface protein, VspH | E258 | None | E258 | None | None |
| BP951000_RS06935 | Hypothetical protein | S9, I10, V13, R298 | None | None | None | S9, I10, V13, R298 |
| BP951000_RS11380 | ToxinA | M126, M154, I278 | None | M126 | M154, I278 | None |
| BP951000_RS03405 | Hypothetical protein | M1, R2, L3, K4, F5, F6, F7, L8, I9, F10, L11, F12, L13, S14, L15, S16, L17, Y18, T19, Q20, D21, N22, E23, A24 | None | None | None | M1, R2, L3, K4, F5, F6, F7, L8, I9, F10, L11, F12, L13, S14, L15, S16, L17, Y18, T19, Q20, D21, N22, E23, A24 |
| BP951000_RS00185^η^ | Hypothetical protein | 58 variations | Not determined^µ^ | Not determined^µ^ | Not determined^µ^ | Not determined^µ^ |
| BP951000_RS10320 | Hypothetical protein | L18, D48, E55, F251, G285, E400, Y401, G402, I403, F404, T405, K406, Q407, L408, A409, I410, S411, F412, I413, P414, I415, N416, I417, R418, F419 | G285 | None | E400, Y401, G402, I403, F404, T405, K406, Q407, L408, A409, I410, S411, F412, I413, P414, I415, N416, I417, R418, F419 | L18, D48, E55, F251 |
| BP951000_RS05445 | DUF3575 domain-containing protein | K2, I7, A79, N87, H89, K158 | K158 | N87, H89 | A79 | K2, I7 |
| BP951000_RS08300 | Tia invasion determinant | L143, N156, S200 | None | None | L143, N156, S200 | None |
| BP951000_RS05490^η^ | Tia invasion determinant | 61 variations | Not determined^µ^ | Not determined^µ^ | Not determined^µ^ | Not determined^µ^ |
| BP951000_RS07500^η^ | Hypothetical protein | 183 variations | Not determined^µ^ | Not determined^µ^ | Not determined^µ^ | Not determined^µ^ |
| BP951000_RS01590 | Hypothetical protein | V205, I215, V221 | None | None | V205, I215, V221 | None |
| BP951000_RS08295 | Tia invasion determinant | N34, I49, V123, S141, I144, V167 | None | N34 | I49, V123, S141, I144, V167 | None |
| BP951000_RS08975 | TonB-dependent receptor domain-containing protein | D32, T371 | None | D32 | T371 | None |
| BP951000_RS06930^η^ | Serpentine receptor domain containing protein | 135 variations | Not determined^µ^ | Not determined^µ^ | Not determined^µ^ | Not determined^µ^ |
| BP951000_RS03290 | Serpentine receptor domain containing protein | None | None | None | None | None |
| BP951000_RS00765^η^ | Serpentine receptor domain containing protein | 70 variations | Not determined^µ^ | Not determined^µ^ | Not determined^µ^ | Not determined^µ^ |
| BP951000_RS01280 | Serpentine receptor domain containing protein | V32, A83, V124, E210, T237 | None | E210 | V32, A83, V124, T237 | None |
| BP951000_RS10445 | Serpentine receptor domain containing protein | G228 | G228 | None | None | None |
| BP951000_RS00365 | Serpentine receptor domain containing protein | V72, Q77, I84, D156, D159, V168, N177, A200, T216, I222, Y226 | Q77, N177, I222, Y226 | D156, D159, A200 | V72, I84, V168, T216 |  |
| BP951000_RS04620 | Serpentine receptor domain containing protein | K2, E95, A140, V194 | None | None | E95, A140, V194 | K2 |
| BP951000_RS04220^η^ | Serpentine receptor domain containing protein | 218 variations | Not determined^µ^ | Not determined^µ^ | Not determined^µ^ | Not determined^µ^ |

^#^Proteins were named as per their annotation in NCBI and Uniprot databases, searched using Protein Accession Number.

ECL – Extracellular loop

ICL – Intracellular loop

TM – Transmembrane

^η^Sequence comparison of these proteins across nine strains of *Brachyspira pilosicoli* revealed variations at more than 40 positions. Therefore, they have been enlisted in tableS4.

^µ^Since sequence variations were present at more than 40 positions, we have not determined whether these variations are present on the TM region or loop region of the predicted proteins.

**Supplementary Table 4:** Sequence comparison across nine strains of *B. pilosicoli* for the proteins having a high number of variations.

| **Locus Identifier** | **Protein name^#^** | **Total variations** |
| --- | --- | --- |
| **GROUP A** |  |  |
| BP951000_RS10215 | Variable surface protein, VspE | F4, V8, T9, V11, F12, V13, L14, A16, S17, D26, N27, T28, L30, F31, I34, N44, V46, T49, G53, T54, V55, F59, K62, A63, N64, T65, G66, L67, T68, Q70, F72, T73, K75, G76, N77, K78, L80, E81, A82, G86, G92, G94, V97, V99, N106, N107, A108, A109, G111, N112, V113, D114, A115, F120, V121, F122, V125, N127, N128, V131, V133, S136, S138, S139, A141, D142, L143, N144, N145, G146, D149, F151, L153, I155, P156, A157, D166, Y170, F173, E174, F175, N176, Q179, S181, K183, E184, G185, T186, V187, N188, Y189, S190, A191, K192, N193, L194, S195, Q197, L198, L200, H201, F202, L203, N204, T205, V206, I207, E208, N209, T211, V212, N213, R217, D219, F220, A221, S222, T230, A231, I232, V233, G234, T235, L236, G237, T241, S242, D243, I244, K245, A246, W247, T248, V249, A250, G251, A252, A253, E254, A255, T256, A257, G258, Q259, E260, R264, Y267, D268, R270, I271, S274, I275, S276, L277, T278, V279, T281, N285, F286, I287, F288, P290, I292, R295, V296, E297, Y299, K301, Q302, G303, G304, K305, L306, E307, S309, V310, Y311, A314, V318, V320, R321, I323, P324, A325, M332, D333, V334, N335, G337, V338, P339, K340, L341, Q342, D343, Q344, S345, T346, T347, P349, I350, A351, G353, N355, S356, S364, N366, K367 |
| BP951000_RS04760 | Variable surface protein, VspD | V4, A8, I9, L11, L12, T13, I14, S16, T17, Y22, N26, S27, D28, I30, D31, V34, D35, A36, F41, V43, R44, M46, V49, D53, M54, I55, R56, V58, V59, V61, R62, D63, E64, S65, E66, T67, S68, F69, A71, F73, D74, S75, T76, T77, Y78, A79, P80, N81, G82, T83, T84, Q86, F87, L88, F90, I91, P92, A96, F102, I104, S106, I107, V109, R114, H116, H117, T118, T119, F120, N121, G122, L123, T124, K125, N126, Y128, G129, L130, S131, E132, L137, T138, M139, T140, M142, D144, S145, I148, I150, V152, Q153, V154, A155, V156, N158, G159, D160, I161, A162, N163, S164, T165, D166, K167, V168, K169, M171, V173, M175, D176, T177, E186, E190, L193, Y194, V195, K196, I199, Q201, L202, E203, N204, T205, E206, V207, A208, N209, S210, K211, E212, K213, A214, E215, S216, F217, G218, D220, F221, A223, Y224, F225, G226, A227, T228, V229, G230, D231, A233, L234, K235, K239, T241, Y242, D243, T244, N247, I252, A253, Y254, S255, Y256, N257, T258, G259, A260, A261, A262, T263, Y264, G265, D266, T267, G269, V270, K271, S272, S273, S274, A275, L276, V277, F278, V279, D280, D281, G282, T283, G284, T285, G286, E288, H289, W292, A293, N295, I296, T299, L300, G301, M302, S303, A304, S306, S310, I311, Y312, V313, A315, L317, G320, A321, V322, T324, Y326, K327, F328, N329, A330, A331, N332, A333, A334, N335, T336, G337, I338, L339, S341, M342, R343, N345, A346, I350, I352, N353, V355, K356, D357, Q359, A364, E365, I366, G367, G369, G370, A371, F372, T373, K374, N375, T376, L377, Q378, N380, G381, A382, T383, T391 |
| **GROUP B** |  |  |
| BP951000_RS09655 | DUF5723 domain-containing protein | K2, K3, L4, L6, L8, T9, A10, L11, C12, I13, T14, A16, Y17, S18, A20, I22, P23, T24, A25, N26, M27, N28, T29, F30, D32, N33, S36, A42, G45, E46, F47, T49, D50, N55, V59, N63, I66, A69, G70, V72, G74, N76, S77, S78, D79, S80, T81, V83, A85, F86, V89, L92, N93, A95, V100, A101, V103, Y104, M106, N107, E108, T109, R110, V111, D112, P115, Y116, S117, A118, T119, S121, Q122, G124, I125, T126, K129, V131, T132, T133, G134, S135, T136, V149, I155, R157, G158, N159, S160, K161, Q162, V163, E165, S169, F170, I171, D172, D173, K174, L176, S177, V178, I179, T182, T183, G184, V189, H190, V192, G195, L196, V197, G199, E200, K202, K204, I205, V207, R208, L209, T210, I211, D213, A214, G215, N216, S217, K220, Q221, G223, L224, T225, K226, T227, D228, V229, A230, N231, Q232, Y233, E235, E236, T237, Q238, T239, G240, Y241, T244, G245, N246, N247, T248, M249, N250, S252, I253, A254, S258, M266, T267, G268, N270, G272, T274, G276, I279, G281, M282, G283, K285, S286, A287, Y288, Y289, S290, T291, Y292, K293, N294, T295, V297, N298, G299, T300, T301, G302, E303, R304, K305, Q306, S307, V309, T310, E313, G314, V315, F316, N319, A320, N323, A324, T327, S331, L332, A333, N335, R336, V337, R338, I340, M341, K344, V345, A348, I349, N350, V351, A354, N356, V357, K358, V360, E361, K362, E364, I365, T366, D367, Q368, T369, L371, E372, T373, T374, T375, S377, G378, A379, K380, S381, S382, L383, V384, Q385, I390, L392, I394, A397, F398, V401, D402, F406, V410, S411, R413, L414, G415, F416, A417, M418, T419, T420, G421, S422, Y423, A425, L426, G428, G429, Y431, K432, T433, F334, N435, Y436, N437, N438, G439, E441, T443, M444, N445, L446, T448, M450, I453, V454, G455, E456, D457, F458, I460, Y463, A465, A466, R467, A468, G469, N470, T471, A472, T473, T474, T475, P476, S477, P478, S479, L480, L481, G482, I483, D484, S485, W486, A488, R493, L494 |
| BP951000_RS08285 | Trep protein | M1, K2, K3, Y4, I5, L6, F7, F8, L9, I10, F11, S12, I13, S14, V15, L16, R205, W207, H240, T305, T322, A349, Q376, R379, E424, R509, V535, K538, T552, T564, S574, A676, L728, V783, A811, S813, S866, I873, A893, R906, S908, I913, S919 |
| BP951000_RS04505 | Variable surface protein VspH | R3, I4, L5, M6, S7, I8, L9, L10, M11, V13, L14, S15, I16, Q17, F19, A20, K23, S24, D25, M27, M30, M37, I39, L44, V46, A48, P50, R51, W53, F55, V56, A57, G61, I64, A65, G67, L68, H71, T73, G74, A75, P76, N77, T78, Q79, Q80, E81, N82, T83, E84, N85, N86, Y87, K88, K89, G90, V91, D92, K93, A98, V100, A101, F102, D105, S106, D107, L108, F109, I111, A112, A113, N118, W119, S121, P122, T123, I130, L131, H132, M133, L136, S139, I146, V150, Q152, K153, Y155, V156, N157, D158, K159, S160, K162, T164, M165, I167, A170, I171, G173, A178, E179, D180, I181, A183, L184, L187, F189, Y190, S196, T197, I198, K199, A200, L201, D202, F203, K204, D205, A206, S207, F208, T209, G215, E216, F217, M219, K222, I223, L224, T225, E226, N227, I228, K229, I230, F234, R237, F238, A240, A243, T244, T245, Y246, K247, N248, I249, D250, E251, A252, N253, R254, G255, S256, I257, L258, S260, Y261, A262, V263, S264, F268, I269, D271, D272, P273, G274, G275, T276, G277, A278, N279, I280, A281, A282, G284, A285, N286, A287, S288, G289, T290, L291, Q292, S301, G302, Y303, K306, E307, L311, L313, I315, V317, T320, A324, E326, S329, F332, L338, I340, V341, E345, T348, F349, G350, Q351, H352,A353, W354, E355, D356, V357, A358, S359, R360, H361, R362, R363, T364, N365, Y368, A369, F370, A376, V383, K384, N385, T390, L392, S397,T398, V399, A400, G401, D402, L403, S404, T405, A406, S407, S408, T410, I411, F413, A415, S416, I419, W421 |
| BP951000_RS00185 | Hypothetical protein | M1, K2, R3, I4, Y5, I6, I7, T8, I9, L10, F11, F12, T13, F14, S15, F16, L17, L18, Y19, S20, Q21, D22, S23, V24, S25, N26, D27, Y28, S29, F30, T31, N32, P33, F34, R35, I36, Y37, D38, V39, D40, K41, Y42, Y43, V44, G45, W46, Q47, D48, P49, R50, A51, F52, I53, G54, R55, L56, M147, P252 |
| BP951000_RS00765 | Serpentine receptor domain containing protein | Q163, M164, Y165, V166, I167, P168, Y169, I170, K171, L172, T173, F174, D175, W176, F177, F178, S179, D180, I181, N182, Y183, K184, T185, R186, E187, F188, V189, D190, T191, R192, D193, L194, G195, I196, G197, F198, Y199, L200, G201, Y202, N203, F204, G205, P206, K207, S208, K209, N210, Y211, I212, G213, T214, D215, S216, F217, D218, I219, G220, L221, Q222, L223, S224, L225, R226, F227, K228, P229, A230, K231, N232 |
| BP951000_RS04220 | Serpentine receptor domain containing protein | K2, F3, V4, K5, Y6, F7, L8, L9, I10, N11, F13, I14, I15, T16, I17, T18, Y19, S20, A21, A23, S24, L25, I27, N28, F29, Q30, G31, H32, Y33, A36, F37, P38, F39, N40, S41, I42, K43, V44, N45, D46, N47, Y48, K49, N50, T51, I52, Y53, D54, S55, V56, D57, G58, T59, L60, F62, E63, G64, T65, L66, F67, F68, Q69, I70, N72, Y73, F74, Q75, L76, F77, E78, D80, Y81, T82, R83, I84, I85, K86, G87, V88, L90, F91, G92, D93, I94, F96, S97, I98, N99, A100, L101, M102, S103, D104, Y105, K106, E107, N108, N109, N110, K111, Y112, T113, E114, V115, L116, G117, F118, Y119, S120, M121, L122, V123, A125, T126, V127, L129, F131, S132, K133, M134, I136, G137, G139, T140, I142, I143, A144, Y147, A148, M149, V150, A151, S152, K154, G156, G157, V158, M159, S160, A161, P162, D163, D164,L165, D166, N167, W168, V170, N171, D172, M173, R174, N175, L176, F177, K178, A179, I181, M182, L187, T188, I189, E190, G191, F192, L193, L195, V196, P197, N198, F199, A200, V201, T202, L203, G205, M207, M208, N210, G212, M213, Q214, Y215, K216, D218, V219, V220, N221, N222, N223, L224, G225, G226, N227, I228, Y229, N230, Q231, Y232, N233, F234, D236, F237, S238, I239, G240, L241, I242, L243, S246, F247, G248, R249, S250, D251, G252, Y253, N254 |
| BP951000_RS06930 | Serpentine receptor domain containing protein | M1, I2, L3, R4, K5, T6, F7, F10, V11, L12, L13, I14, L15, F18, N19, L21, A22, N23, K25, F26, I28, S29, P30, R31, L32, A34, D35, I36, N37, I38, A39, E40, T41, K42, N43, Y44, N45, S46, L47, L48, L49, D50, L53, I55, F57, S58, I59, G62, K64, L65, N67, I68, N69, L70, L71, S72, G73, I74, D75, F80, N83, E84, H85, M86, L87, I89, V90, D91, K92, T94, S95, L96, R99, V100, T101, F103, G104, V105, Y106, A107, V109, D114, L115, S116, K117, N118, N119, Q120, N121, L123, I124, I126, Q131, N133, P134, L135, N139, Y140, K141, S142, R150, S152, R155, K156, F157, F158, E159, T160, K162, T163, D164, L166, A167, V169, H170, N175, N179, Y180, N181, M183, N188, Y189, H190, D191, V192, K193, Y195, L200, F202, F204, S205, F206, F207, E209, N210 |
| BP951000_RS05490 | Tia invasion determinant | M1, K2, V3, Y4, A8, V9, T14, F17, A21, T23, E24, E26, K28, K29, N30, K31, V32, L33, G35, I36, Y46, T51, F65, E71, N72, V79, N87, D89, F95, I99, T100, S103, K104, V106, T108, E110, A114, A117, A121, G129, T132, D133, H134, E140, L142, W144, K158, V163, D172, I173, Y174, A177, K178, R183, L185, M187, V188, M195, V197, I201, W202 |
| BP951000_RS03215 | TonB-dependent siderophore receptor | M1, K2, F4, I5, Y7, L9, F10, A11, S12, I13, M15, T20, K35, V36, N41, T42, Q43, T44, N45, N46, E47, Y48, S49, N50, V51, S52, K53, I55, N58, V59, N61, K62, T63, E70, N75, K78, I80, E82, A85, N92, G104, S105, A106, K141, S147, Q258, E285, V291, F306, K307, E308, L332, E342, D377, S483, A506, I569, Y591, I593 |
| BP951000_RS07500 | Hypothetical protein | M1, K2, K3, I4, I5, V6, L7, G8, L9, M10, L11, I12, S13, S14, S15, L16, V17, Y18, S19, H20, S21, L22, G23, I24, G25, M26, Y27, I28, P29, L30, G31, G32, S33, L34, P35, S36, F37, Y38, S39, D40, N41, A42, E43, A44, T45, S46, F47, F48, S49, P50, K51, S52, A53, F54, E55, V56, G57, V58, I59, F60, N61, P62, R63, V64, N65, F66, N67, I68, G69, D70, G71, T72, H73, T74, V75, S76, L77, G78, V79, D80, V81, G82, W83, Y84, R85, D86, A87, F88, K89, F90, A91, S92, S93, T94, D95, V96, T97, H98, E99, F100, D101, T102, V103, M104, T105, G106, L107, N108, L109, E110, W111, R112, P113, L114, L115, F116, Q117, L118, G119, V120, G121, G122, G123, V124, K125, F126, P127, F128, L129, G130, K131, Y132, I133, E134, G135, N136, N137, K138, M139, A140, L141, S142, G143, G144, A145, F146, A147, S148, R149, Y150, N151, N152, V153, F154, I155, P156, Y157, I158, R159, L160, Y161, T162, G163, I164, N165, I166, I167, F168, I169, S170, L171, S172, L173, Y174, V175, N176, F177, D178, I179, P180, Y181, L182, Q183 |

^#^ Proteins were named as per their annotation in NCBI and Uniprot databases, searched using Protein Accession Number.

**Supplementary Table 5:** BlastP output of all identified OMBBs of *Brachyspira pilosicoli* against mentioned species of *Brachyspira* below in the table.

| **Identified OMBBs (*B. pilosicoli*)** | **Pathogenic species** | | | | **Non-pathogenic species** | |
| --- | --- | --- | --- | --- | --- | --- |
|  | ***B. aalborgi*** | ***B. hyodysenteriae*** | ***B. intermedia*** | ***B. alvinipulli*** | ***B. murdochii*** | ***B. innocens*** |
| **Group A** |  |  |  |  |  |  |
| WP_013244106.1 | **+** | **+** | **+** | **+** | **+** | **+** |
| WP_013244995.1 | **-** | **+** | **+** | **-** | **+** | **+** |
| WP_013243917.1 | **-** | **+** | **+** | **-** | **+** | **+** |
| WP_013243377.1 | **+** | **+** | **+** | **+** | **+** | **+** |
| WP_013243376.1 | **+** | **+** | **+** | **+** | **+** | **+** |
| WP_013244459.1 | **+** | **+** | **+** | **+** | **+** | **+** |
| WP_013242998.1 | **-** | **+** | **+** | **+** | **+** | **+** |
| WP_013243940.1 | **-** | **+** | **+** | **+** | **+** | **+** |
| WP_013243193.1 | **+** | **+** | **+** | **+** | **+** | **+** |
| WP_013243655.1 | **+** | **+** | **+** | **+** | **+** | **+** |
| WP_013244081.1 | **+** | **+** | **+** | **+** | **+** | **+** |
| WP_013244750.1 | **+** | **+** | **+** | **+** | **+** | **+** |
| WP_041747714.1 | **+** | **+** | **+** | **+** | **+** | **+** |
| **Group B** |  |  |  |  |  |  |
| WP_041747843.1 | **+** | **+** | **+** | **+** | **+** | **+** |
| WP_013244879.1 | **+** | **-** | **+** | **-** | **-** | **+** |
| WP_013244607.1 | **+** | **+** | **-** | **+** | **+** | **+** |
| WP_015274839.1 | **+** | **-** | **+** | **-** | **+** | **-** |
| WP_013243854.1 | **-** | **+** | **+** | **+** | **+** | **+** |
| WP_013243861.1 | **-** | **+** | **+** | **+** | **+** | **+** |
| WP_013242999.1 | **+** | **-** | **+** | **+** | **+** | **+** |
| WP_013244339.1 | **+** | **+** | **+** | **+** | **+** | **+** |
| WP_013243647.1 | **+** | **-** | **+** | **+** | **+** | **+** |
| WP_013244641.1 | **+** | **+** | **+** | **+** | **+** | **+** |
| WP_041747935.1 | **+** | **+** | **+** | **+** | **+** | **+** |
| WP_013243225.1 | **+** | **+** | **+** | **+** | **+** | **+** |
| WP_013243896.1 | **-** | **+** | **+** | **-** | **+** | **+** |
| WP_014933009.1 | **-** | **+** | **+** | **-** | **+** | **+** |
| WP_014936494.1 | **+** | **+** | **+** | **+** | **+** | **+** |
| WP_013243815.1 | **+** | **+** | **+** | **+** | **+** | **+** |
| WP_013245039.1 | **+** | **+** | **+** | **+** | **+** | **+** |
| WP_013243037.1 | **+** | **-** | **+** | **+** | **+** | **+** |
| WP_013244338.1 | **+** | **+** | **+** | **+** | **+** | **+** |
| WP_013244050.1 | **+** | **+** | **+** | **+** | **+** | **+** |
| WP_013244610.1 | **+** | **+** | **+** | **+** | **+** | **+** |
| WP_228369485.1 | **+** | **+** | **+** | **+** | **+** | **+** |
| WP_013244059.1 | **+** | **+** | **+** | **+** | **+** | **+** |
| WP_041747581.1 | **-** | **+** | **+** | **+** | **+** | **+** |
| WP_041747873.1 | **+** | **+** | **+** | **+** | **+** | **+** |
| WP_187287137.1 | **-** | **+** | **+** | **+** | **+** | **+** |
| WP_181893515.1 | **+** | **+** | **+** | **+** | **+** | **+** |
| WP_013244745.1 | **-** | **+** | **+** | **+** | **+** | **+** |
| WP_013243185.1 | **+** | **+** | **+** | **+** | **+** | **+** |
| WP_013243063.1 | **+** | **-** | **+** | **+** | **+** | **-** |

**Supplementary Figures**

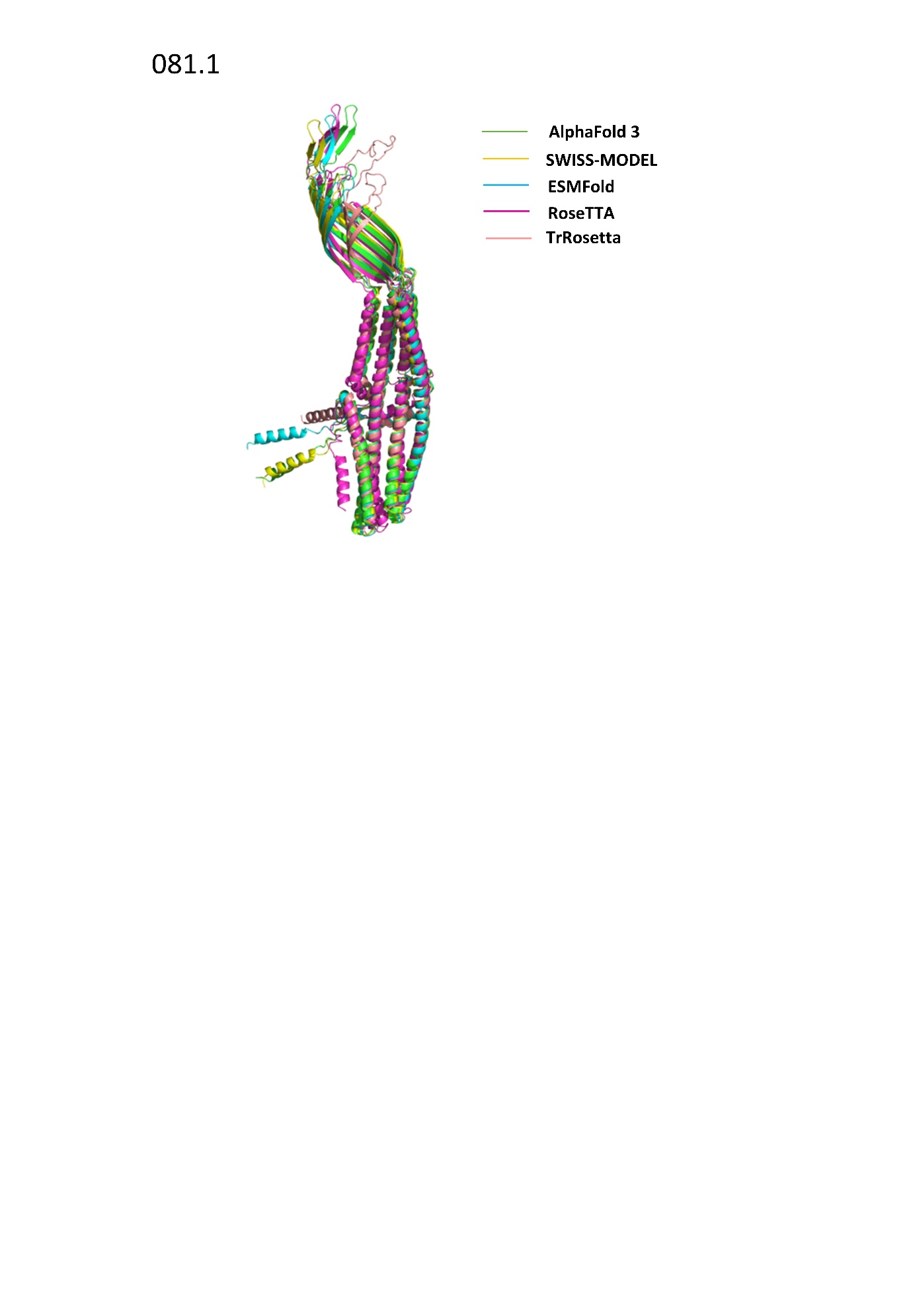
**Figure S1:** Structural alignment of monomeric structures of BP951000_RS05600 generated by different modelling tools (AlphaFold 3, ESMFold, RoseTTA, SWISS-MODEL and TrRosetta) using US-Align server resulted an RMSD of 3.96Å.

**Figure S2:** Structural alignment of monomeric structures of BP951000_RS09000 generated by different modelling tools (AlphaFold 3, ESMFold, RoseTTA, SWISS-MODEL and TrRosetta) using US-Align server resulted an RMSD of 3.85Å.

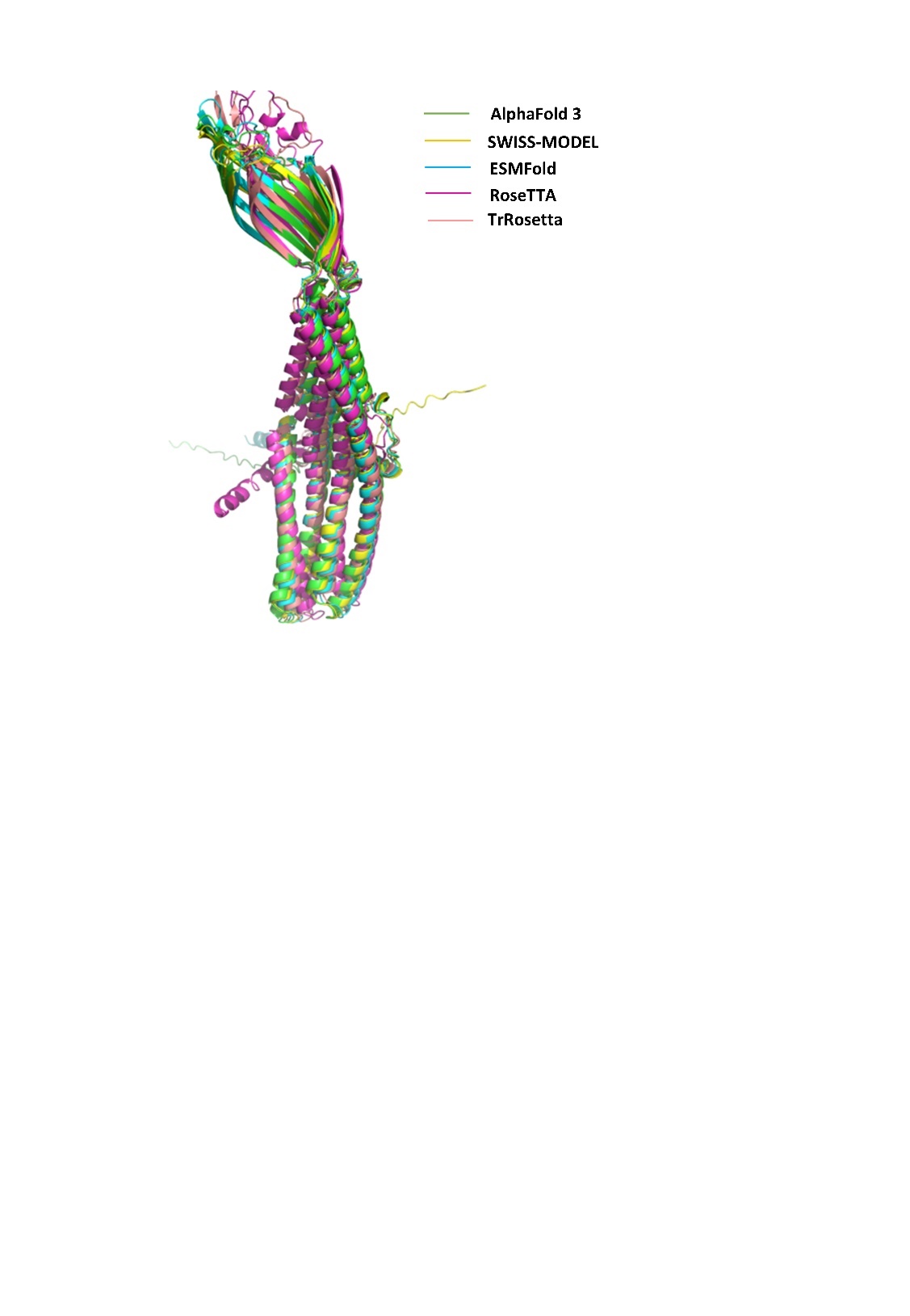

**Figure S3:** Structural alignment of monomeric structures of BP951000_RS06235 generated by different modelling tools (AlphaFold 3, ESMFold, RoseTTA, SWISS-MODEL and TrRosetta) using US-Align server resulted an RMSD of 3.26Å.

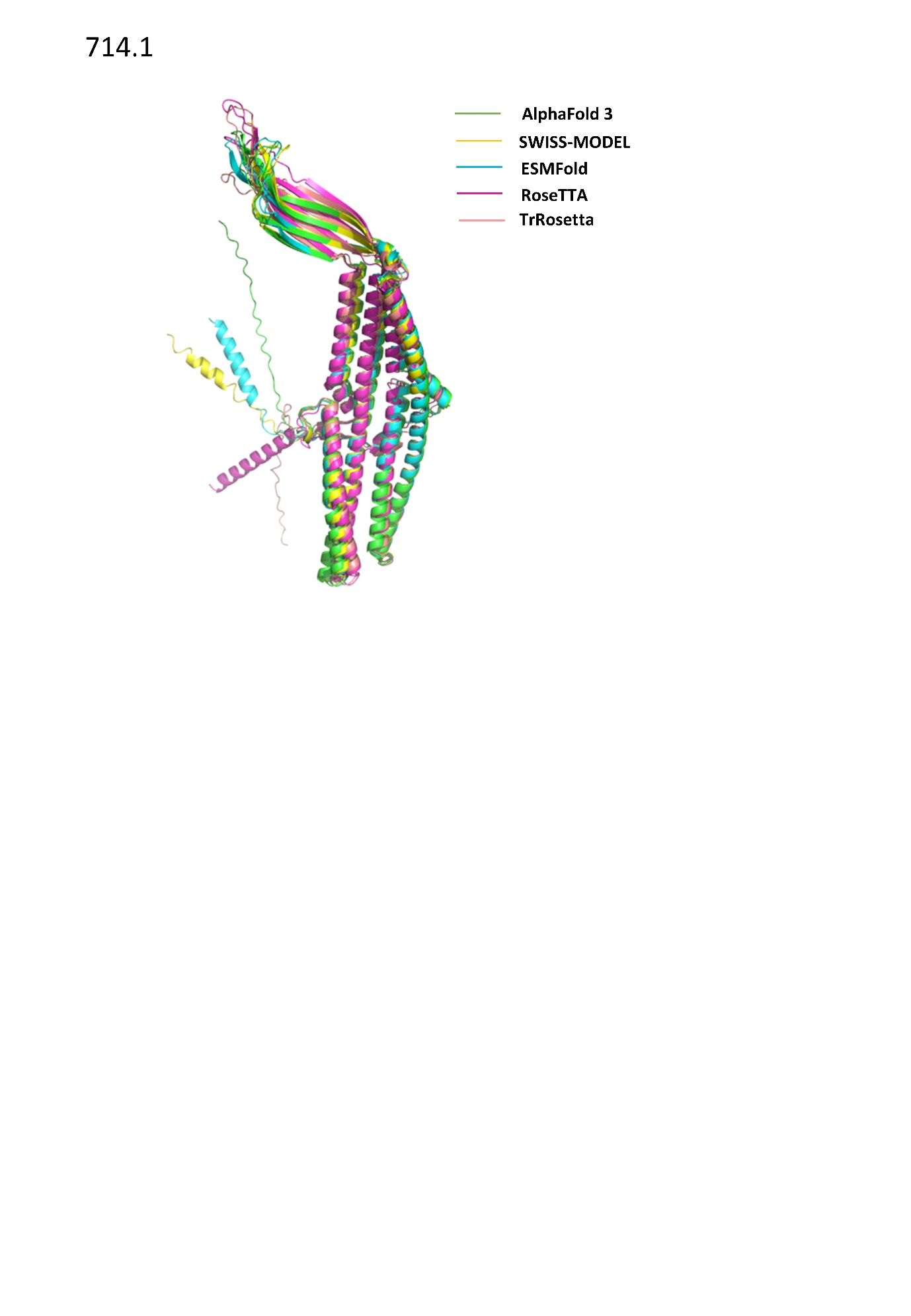

**Figure S4:** Structural models of the Trep protein of *B. pilosicoli*, *Bordetella pertussis* and *E. coli*.
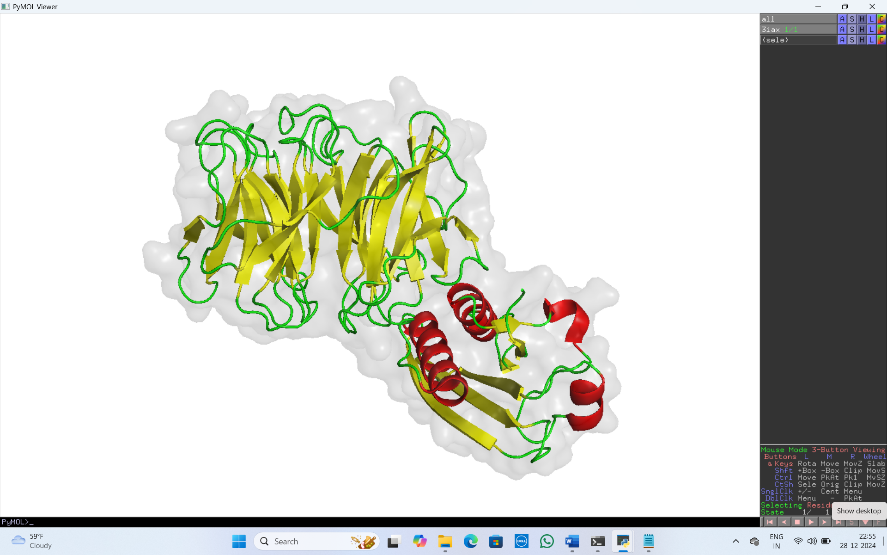

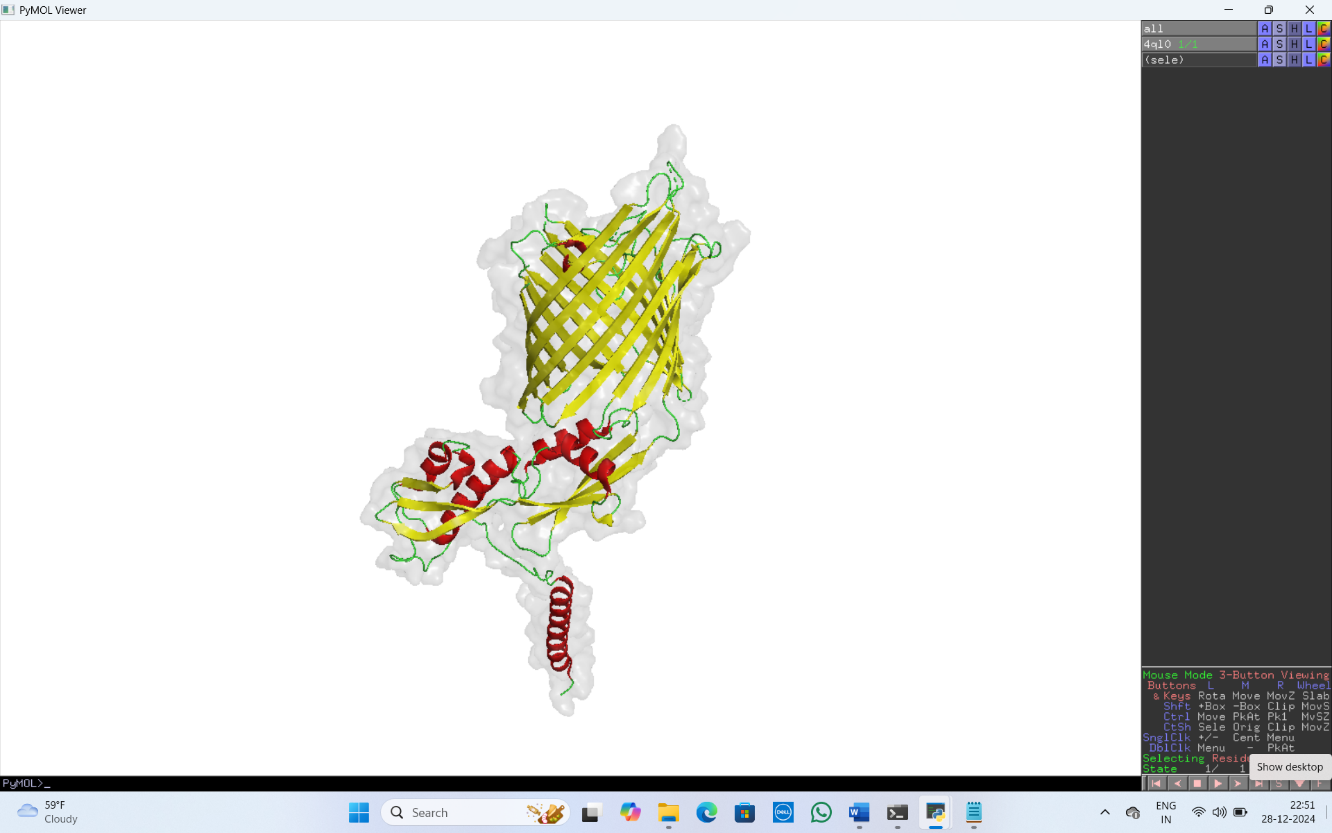

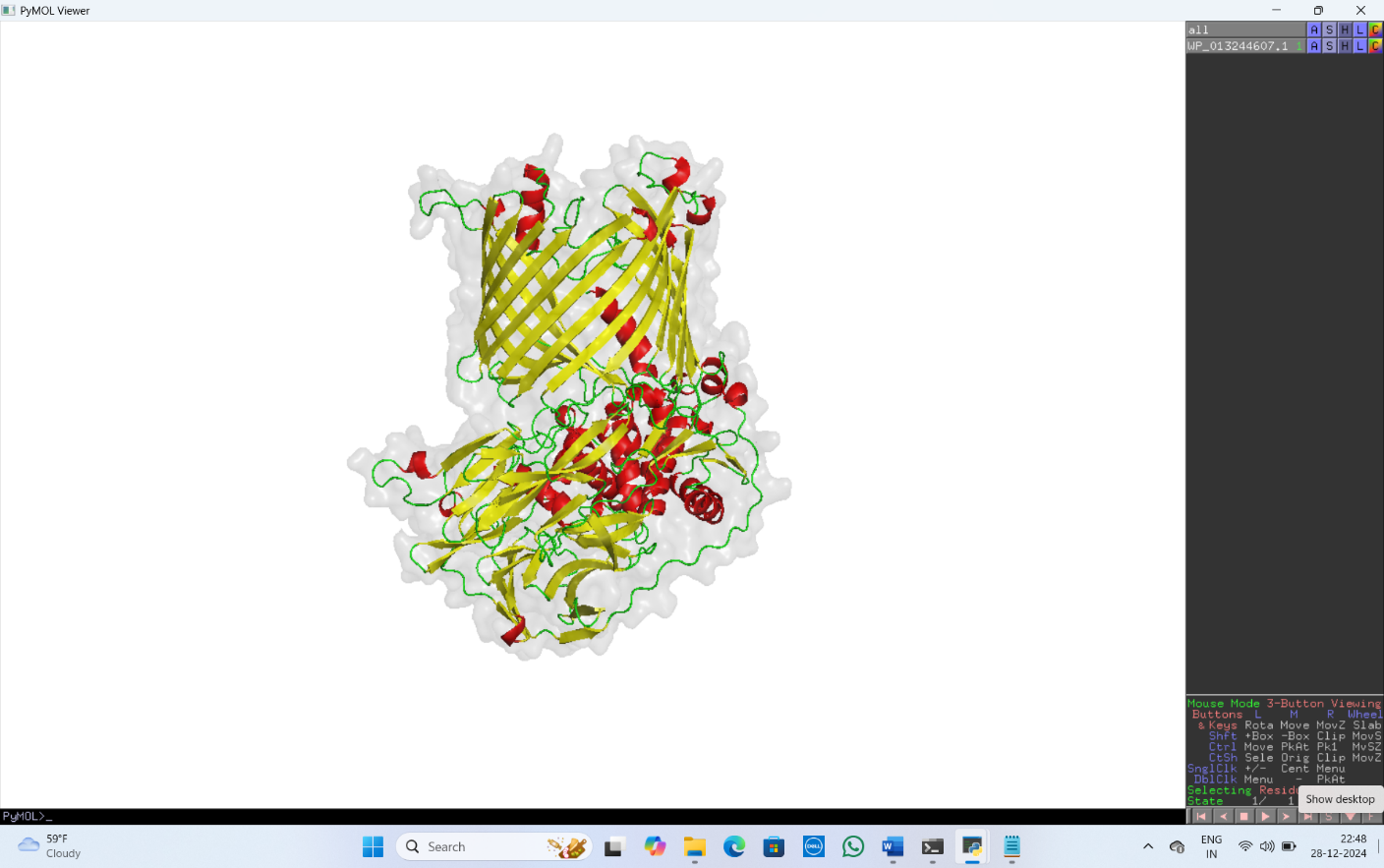

TolB of *E. coli* K-12 (PDB ID: 3IAX)

FhaC of *Bordetella pertussis* (PDB ID: 4QL0)

BP951000_RS08285, Trep protein of *B. pilosicoli*

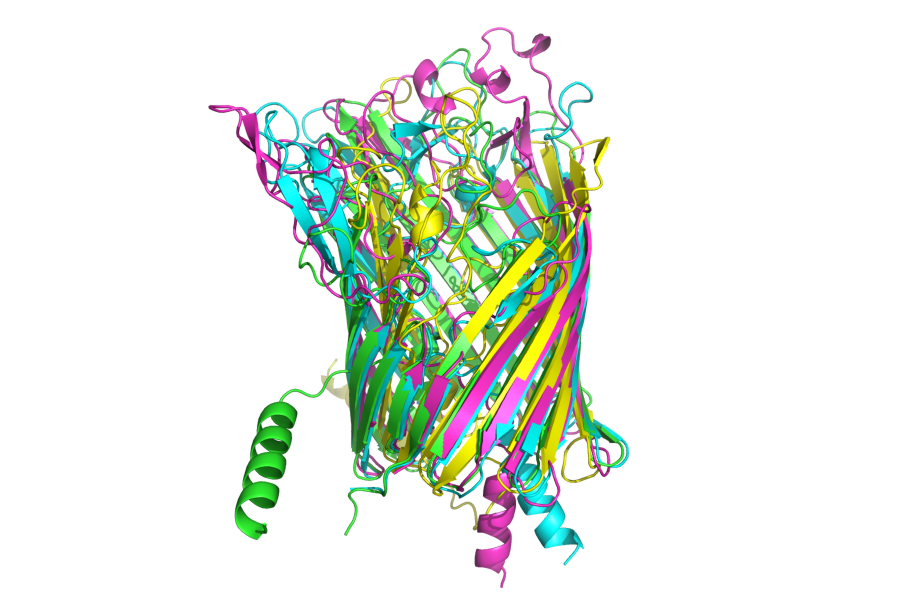
**Figure S5**: Structural alignment of BP951000_RS04760, BP951000_RS10215, BP951000_RS04505 and BP951000_RS01090 using US-align server yielded an RMSD of 4 Å. Cyan color represents BP951000_RS04760 (VspD), green color represents BP951000_RS10215 (VspE), magenta color represents BP951000_RS04505 (VspH) and yellow color represents BP951000_RS01090 (VspH, 14-stranded β-barrel protein).
